# Olfactory Projection Neurons From the Moth Antennal Lobe Lateral Cluster exhibit Diverse Morphological and Neurophysiological Characteristics

**DOI:** 10.1101/274506

**Authors:** Seong-Gyu Lee, Christine Fogarty Celestino, Jeffrey Stagg, Christoph Kleineidam, Neil J. Vickers

**Affiliations:** Shin-Hwa ED Tech Co. Ltd., Ulsan Technopark Rm. A-409, Ulsan, South Korea; Science Department, Juan Diego High School, 300 East 11800 South, Draper, UT 84020; Texas Tech University Health Sciences Center El Paso, 4800 Alberta Avenue, El Paso, TX 79905; Neurobiologie, University of Konstanz, Universitätsstraße 10, D-78457 Konstanz, Germany

## Abstract

Olfactory projection neurons convey information from the insect antennal lobe (AL) to higher centers in the brain. Many studies on moths have reported excitatory projection neurons with cell bodies in the medial cell cluster (*mc*PNs) that predominantly send an axon from the AL to calyces of the mushroom body (CA) via the medial antennal lobe tract (mALT) and then to the lateral horn (LH) of the protocerebrum. These neurons tend to have dendritic arbors restricted to a single glomerulus (i.e. they are uniglomerular). In this study, we report on the physiological and morphological properties of a group of pheromone-responsive olfactory projection neurons with cell bodies in the moth AL lateral cell cluster (*lc*PNs) of two heliothine moth species. While *mc*PNs typically exhibit a narrow odor tuning range related to the restriction of their dendritic arbors within a single glomerulus, *lc*PNs exhibited an array of morphological and physiological configurations. Pheromone-responsive *lc*PNs varied in their associations with glomeruli (uniglomerular and multiglomerular), dendritic arborization structure and connections to higher brain centers with projections primarily through the lateral antennal lobe tract and to a lesser extent the mediolateral antennal lobe tract to a variety of protocerebral targets including ventrolateral and superior neuropils as well as LH. Physiological characterization of *lc*PNs also revealed a diversity of response profiles including those either enhanced by or reliant upon presentation of a pheromone blend. These responses manifested themselves as higher maximum firing rates and/or improved temporal resolution of pulsatile stimuli. *lc*PNs therefore participate in conveying a variety of olfactory information relating to qualitative and temporal facets of the pheromone stimulus to a more expansive number of protocerebral targets than their *mc*PN counterparts. The role of *lc*PNs in the overall scheme of olfactory processing is discussed.

## Introduction

Olfactory sensory neurons (OSNs) located in cuticular hairs covering insect antennae, called sensilla, project axons into the brain and form arbors restricted to olfactory glomeruli in the antennal lobe (AL). OSNs form synaptic connections in the AL with local interneurons (LNs) and projection neurons (PNs) that relay olfactory information from the AL to the protocerebrum. In the large sphingid hawkmoth, *Manduca sexta*, initial studies revealed the cell bodies of AL neurons to be associated with one of 3 different clusters located at the margin of the AL neuropil: a medio-dorsal cluster (*mc*), a lateral cluster (*lc*) and a small antero-ventral cluster (*ac*) (Matsumoto & Hildebrand 1981; Homberg et al. 1988). A detailed study by Homberg et al. (1988) described the anatomy of antenno-cerebral pathways in *M. sexta*, and revealed three primary tracts (names conform to the current *Drosophila* brain map, Ito et al., 2014, with corresponding names in parentheses from Homberg et al., 1988): medial antennal lobe tract, mALT (formerly inner antennocerebral tract, iACT); medio-lateral ALT, mlALT (formerly medial ACT, mACT) and lateral ALT, lALT (formerly outer ACT, oACT). Connections through PNs with cell bodies in each of the clusters contributed to each of 3 primary tracts.

Investigations of PNs in many different moth species have included evaluation of neurophysiological responses to stimulation with odor often partnered with morphological characterization of individual neurons through staining of their antennal lobe dendritic arborizations and terminal target arbors in the protocerebrum (Christensen and Hildebrand, 1987; Kanzaki et al., 1989, 2003; Christensen et al., 1991; Hansson et al., 1991; 1994; Anton and Hansson, 1995, 1997; Lei and Hansson, 1999; Vickers et al., 1998; Lei and Hansson, 1999; Heinbockel et al., 2004; Reisenman et al., 2004; Zhao and Berg, 2010; Zhao et al., 2014). Male moth PNs sensitive to compounds predominantly found in female pheromone blends have dendritic arborizations in the sexually-dimorphic cluster of glomeruli known as the macroglomerular complex (MGC) of the AL (Matsumoto and Hildebrand, 1981; Homberg et al., 1988; Hildebrand 1996). Analysis of fiber counts in output tracts suggested that MGC-PNs primarily exited the AL via the mALT and lALT (Homberg et al., 1988). A further morphological study of Rø et al. (2007) confirmed many of the earlier observations of Homberg et al. (1988) while also advancing our understanding of the protocerebral targets of PNs associated with the different AL output pathways. This work focused on results from female *Heliothis virescens* and, as a consequence, featured PNs with dendritic arborizations in sexually-isomorphic or ordinary glomeruli.

Many studies in different moth species have demonstrated that MGC-PNs typically exhibited uniglomerular dendritic arborizations, responded primarily to single pheromonal odorants, had *mc*-associated somata and projected through the mALT (herein denoted m-*mc*PNs) first to the calyces of the mushroom body (CA) before terminating in the lateral neuropils of the medial protocerebrum such as the lateral horn (LH) (e.g. Kanzaki et al., 1989, 2003; Christensen et al., 1991; Anton et al., 1997; Vickers et al., 1998; Kanzaki et al., 2003; Vickers and Christensen, 2003; Lee et al., 2016). Neurons with somata located within the *lc* tended to be intrinsic inhibitory local interneurons (iLNs) that did not extend an axon beyond the AL (Christensen et al., 1993). However, even though anatomical studies clearly indicated that the *lc* also housed the somata of PNs that convey information to a variety of higher brain centers (Homberg et al., 1988; Rø et al., 2007; Ian et al., 2016a,b), our understanding of their function in moth olfactory processing is extremely limited primarily because we have little information regarding the neurophysiological response profiles of these neurons to olfactory stimulation.

Studies and species in which the paired morphological and neurophysiological data for the occasional *lc*PN has been reported include: *O. nubilalis* (Anton et al., 1997; Kárpáti et al., 2008, 2010); *A. segetum* (Hansson et al., 1994; Wu et al., 1996); female *S. littoralis* (female: Anton and Hansson, 1994; male: Anton and Hansson, 1995); *H. virescens, H. zea, H. subflexa* (Christensen et al., 1989; Vickers et al., 1998; Vickers and Christensen, 2003; Løfaldi et al., 2012), *H.virescens*/*H.subflexa* hybrids (Vickers, 2006); *H. virescens*/*H. zea* imaginal disc transplants (Vickers et al., 2005); *Manduca sexta* (males: Heinbockel et al., 1998, 2004; females: Reisenman et al., 2004) and *Trichoplusia ni* (Anton and Hansson, 1999). (Christensen et al., 1991; Anton and Hansson, 1995; Vickers et al., 1998, 2003; Kanzaki et al., 2003; Vickers and Christensen, 2003; Vickers 2006). These studies suggest that *lc*PNs are morphologically distinct from their m-*mc*PN counterparts both in terms of their glomerular associations and the pathways followed to protocerebral centers. Several examples exist of *lc*PNs that exhibited multiglomerular dendritic arborizations and this difference from most m-*mc*PNs subtends distinct response profiles. In *Bombyx mori* reported stains of six *lc*PNs in which these neurons had multiglomerular MGC dendritic arbors (cumulus and toroid compartments) and projected through either the mlALT (ml-*lc*PNs) or the lALT (l-*lc*PNs) to an area identified as the inferior lateral protocerebrum (ILPC) (Kanzaki et al., 2003). Intracellular recordings were made from only one *lc*PN that responded to the pheromone bombykol and less vigorously to a behavioral antagonist, bombykal. Stimulation with a 9:1 blend of these two components did not appear to elicit a response that differed from bombykol alone (Kanzaki et al., 2003). Nitric oxide-induced anti-cGMP immunohistochemistry was used in an ensuing study (Seki et al., 2005) and revealed a pyramid-shaped region of the ILPC (ΔILPC). The ILPC was not recognized as a discrete neuropil in the recent detailed reconstructions of the *Drosophila* brain but was redefined parts of several developmentally distinct neuropils including the lateral part of the superior clamp (SCL), the inferior part of the superior lateral protocerebrum (SLP) as well as small superior-most parts of two ventrolateral protocerebral neuropils (VLPN) – the posterior ventrolateral protocerebrum (PVLP) and posterior lateral protocerebrum (PLP) (Ito et al., 2014). Since the ΔILPC region is now recognized to incorporate parts of both inferior and superior neuropils, in the current study we refer to this area as the ΔLP (Δ-lateral protocerebrum). Intracellular staining of m-*mc*PNs with Lucifer yellow revealed details of their projection patterns first in mushroom body calyx (CA) and then in ΔLP. A spatial separation of m-*mc*PN terminal arbors according to association with single MGC glomeruli was noted both in the mushroom body calyx and ΔLP (Seki et al. 2005). Unfortunately, no individual l-*lc*PNs were stained (Seki et al., 2005) although re-evaluations of l-*lc*PNs stained in the previous study (Kanzaki et al., 2003) revealed that they too arborized in the ΔLP. A further study revealed details of a protocerebral pathway for processing pheromonal information in *B. mori* (Namiki et al. 2014). In addition to input connections from the AL through either mALT via CA or directly through mlALT/lALT into ΔLP, a significant output connection between ΔLP and another neuropil, the superior medial protocerebrum (SMP), was characterized. In *Heliothis virescens*, an *lc*PN with a glomerular dendritic arbor restricted to the cumulus compartment of the MGC responded vigorously to blends of behaviorally active odorants but not to constitutive odorants presented singly (Fig. 11 in Vickers et al., 1998b).

Based on previous reports of lcPNs, Baker and Hansson (2016) proposed that *lc*PNs are ‘chromatic’ with respect to central processing in that they convey information to the protocerebrum about the presence of a pheromone blend. They advanced that chromatic PNs had poorer temporal resolution of pulsatile stimuli compared to m-*mc*PNs and postulated a role in activating circuits that drive longer-lasting moth flight behaviors such as counterturning. Heinbockel et al. (2004) reported that a type of antennal lobe PN in *M. sexta* males belonging to a physiological category that was excited by bombykal (BAL^+^) and inhibited by a (E,Z)-11,13-pentadecadienal (C15^−^). A single stained PN in this BAL^+^/ C15^−^ group of 12 recordings revealed an l-*lc*PN with dendritic arbors restricted to a single MGC glomerulus and an unusual projection to the lateral protocerebrum. In comparison to BAL^+^-specialist PNs, these neurons exhibited improved tracking of stimulus intermittency. Nonetheless, despite the broad array of species in which *lc*PNs have been reported, a systematic evaluation of both neurophysiological and morphological properties of individual neurons is lacking for even a single moth species.

In this study, we report that *lc*PNs in *H. virescens* and *H.subflexa* males with can be categorized into diverse morpho-neurophysiological groups with respect to their responses to individual pheromone components and behaviorally relevant blends (Teal et al., 1981, 1986; Vetter and Baker, 1983; Vickers 2002; Vickers and Baker 1997, Vickers et al., 1991). These neurons distribute a spectrum of olfactory information to a large number of central brain areas suggesting that they subtend an important role in the protocerebral processing of olfactory information and its integration with outputs from other sensory structures.

## Materials and Methods

### Insects

*H. virescens* larvae were fed on a modified pinto bean diet (Shorey and Hale, 1965). *H. subflexa* males were corn-soy blend diet (A. Sheck, personal communication). Males were identified and separated from females at the pupal stage and kept in an environmental chamber on a 14:10 L:D reversed light cycle. Adults were provided with a 10% sucrose solution. Experiments were performed on adult males within 3-5 days of emergence.

### Pheromone

All synthetic pheromone components were acquired from either Sigma-Aldrich (St. Louis, MO) or Bedoukian Research (Danbury, CT) and >95% pure as determined by gas chromatography. Dilutions of individual pheromone components were made in HPLC-grade hexane. Single pheromonal odorants tested (Z11-16:Ald, Z9-14:Ald, Z9-16:Ald, Z11-16:OH, and Z11-16:OAc) were loaded at either 10 or 100ng, as well as 0.05ng (Z9-14:Ald). Hexane alone was used as the control. Depending on the preparation, one or more of the following blends were also tested:

Z11-16:Ald, Z9-14:Ald (1:0.05 or 0.5) (Hv 2mix),

Z11-16:Ald, Z9-14:Ald, Z11-16:OAc, Z11-16:OH (1:0.05:0.1:0.1) (Hv antagonist)

Z11-16:Ald, Z9-16:Ald, Z11-16:OH, Z11-16:OAc (1:0.5:0.1:0.1) (Hs 4mix)

Z11-16:Ald, Z9-14:Ald, Z9-16:Ald, Z11-16:OH, Z11-16:OAc (1:1:1:1:1) (5mix)

Z11-16:Ald, Z9-14:Ald, Z9-16:Ald, Z7-16:Ald, 16:Ald, 14:Ald (1:0.1:0.01:0.01:0.5:0.05) (Hv 6mix)

Z9-16:Ald, Z7-16:Ald, 16:Ald, 14:Ald (1:1:1:1) (Hv minor)

Components and blends were presented as a series of 5 x 40ms pulses (Vickers et al., 1998). Each odorant or blend was loaded onto a 3.5 x 0.5mm filter paper (Whatman) and hexane solvent was allowed to evaporate before insertion into a cartridge (made from a shortened 1mL syringe). Cartridges were capped at both ends, placed within a glass vial and stored at –20° C between experiments.

### Electrophysiology

Intracellular recordings were made using sharp electrodes as described previously for heliothine moths (Christensen et al., 1991, 1995; Vickers et al., 1998). Briefly, the male moth was restrained in a shortened pipette tip and secured with dental wax for dissection. The cuticle and musculature were removed from the anterior portion of the head, exposing the brain. After desheathing the AL, the preparation was placed on the electrophysiology rig where it was bathed with physiological saline and sucrose. Microelectrodes for physiology were pulled (Sutter Instruments) and the electrode tip was backfilled with one of two neuronal tracers: either 3% Lucifer Yellow CH (Sigma-Aldrich, St. Louis, MO) in 0.2M LiCl (2M LiCl backfill) or 5% Neurobiotin™ (Vector Lab, Burlingame, CA) in 0.25M KCl (3M KCl backfill). Resulting electrodes had a resistance of 100-400 MΩ. The microelectrode was lowered into the exposed AL of the moth until stable contact was made with a cell.

Odor cartridges were positioned such that stimulation occurred over the antenna ipsilateral to the recording site. A stimulator controlled a solenoid valve that, when activated, diverted filtered humidified air through a single odor cartridge and over the antenna. Odor stimulation consisted of five pulses of odorant with duration of 40-60ms and an interpulse interval of 300ms.

Neuronal tracers were injected intracellularly with negative current (∼-3nA) for Lucifer Yellow or positive current (∼ 5nA) for Neurobiotin. After removal and fixation overnight in 2.5% formaldehyde, 3% sucrose in 0.1M phosphate buffer, brains that were stained with Neurobiotin (Anton et al., 2003) were then incubated in a blocking solution (1% bovine serum albumin, 0.5 % Triton X-100 in 0.2 M phosphate buffer solution) overnight, followed by 1:80 Streptavidin-Alexa Fluor 555 (Invitrogen, Grand Island, NY) diluted in blocking solution for 12hrs at 4°C and for 6hrs at RT. After rinsing, both LY and Neurobiotin brains were dehydrated in an ethanol series and cleared with methyl salicylate. In some instances, multiple individual neurons were penetrated sequentially with a Neurobiotin-filled microelectrode was injected into each cell that was contacted. Preparations were fixed and cleared as describe above.

### Mass staining

Tungsten needles were created by electrolytically sharpening tungsten rods (0.127mm diameter) in 10% w/v KOH (Brady, 1965). The tips of sharpened needles were then immersed in a slide well containing a small volume of dextran tetramethylrhodamine (TMR) (3000MW – Molecular Probes, Eugene, OR) dissolved in distilled water. As the water evaporated rhodamine crystals were deposited along the needle tip. Prepared needles were stored at RT and set aside for future use. Dissections were prepared as described above. A needle was selected and placed in a pin holder (FST). The needle was inserted directly through the neural sheath into the AL (without need to de-sheath – thereby preserving the morphological integrity of the AL) into the vicinity of the AL where *lc*PNs send primary processes from their cell bodies into the AL neuropil. Preparations were left for 1-2h in a darkened location to allow cells to transport the dye before the brain was removed, fixed and cleared as described above.

### Data Analysis

Cells were monitored on an oscilloscope and recorded on a PCM VHS VCR recorder (A.R. Vetter, Rebersberg, PA). Recordings of interest were subsequently digitized using DataPac 2000 software (Run Technologies, Laguna Niguel, CA) at a sampling rate of 10kHz. Neural activity during stimulation was digitized for 4 seconds following the start of stimulation, along with the 2 seconds prior to stimulation. Action potentials were identified using the “Select Events” function in DataPac by manually setting a voltage threshold for identification of action potentials. Instantaneous spiking frequencies were calculated and exported in a spreadsheet format for graphing and analysis.

Analysis of various response features was performed in order to compare between presentation of a single odorant and a mixture containing this odorant. For practical purposes only neurons in which responses to Z11-16:Ald were elicited could be used because these neurons were also exposed to mixtures in which the concentration of Z11-16:Ald was the same. This criterion was true for 9 of the 12 neuronal physiologies documented in the current study since the two iPNs that responded only to blend stimuli were also excluded. For each volley of action potentials elicited by an odor pulse the latency to onset of the response and peak instantaneous frequency were extracted. Means were calculated for each neuron. A paired t-test was used to compare means across the treatments (Z11-16:Ald/blend) and significance was determined at P<0.05.

### Imaging

Imaging was performed on a Zeiss confocal microscope (LSM 510, Carl Zeiss, Jena, Germany) using the following filter and beam splitter settings: Lucifer Yellow stains: 458nm Argon laser, LP475, NFT490, HFT 458/514, and a blue reflector; Streptavidin-Alexa 555 stains: 543nm HeNe laser, 554∼619 nm band filter. Wholemount brains were imaged with a Plan-Neofluor 20X objective (NA 0.5) in Z-stacks of 1-6μm thick optical sections using LSM software. Following wholemount imaging, select brains were mounted in Spurrs plastic and sectioned on a microtome at 20μm. Regions of interest were then further imaged with a Plan-Neofluor 40X objective (NA 0.75) in Z-stacks of 1μm optical sections.

Post-hoc image adjustments were made for brightness and contrast (Zeiss LSM Image Examiner Version 4.2) with structure labels and scale bars added using Adobe Illustrator. No substantive alterations were made to the images themselves.

### Nomenclature

We adhere to the terminology and acronyms for insect brain regions proposed by Ito et al. (2014) based on their detailed reconstruction of the *D. melanogaster* brain and identify brain regions in the moth accordingly.

## Results

### Antennal Lobe Output Tracts and Protocerebral Targets of lateral cluster Projection Neurons

Results from dextran TMR mass staining (Fig. 1) and indiscriminate intracellular neurobiotin staining (Fig. 2) provided insights into the general anatomical characteristics of moth *lc*PNs. Mass staining primarily of cell bodies in the AL lateral cell cluster with TMR was successful in 7 preparations. Although the degree of staining was variable, the results were consistent across all individuals (Figure 1). *lc*PNs contributed primarily to two AL output pathways: mlALT and lALT.

**Figure 1.**
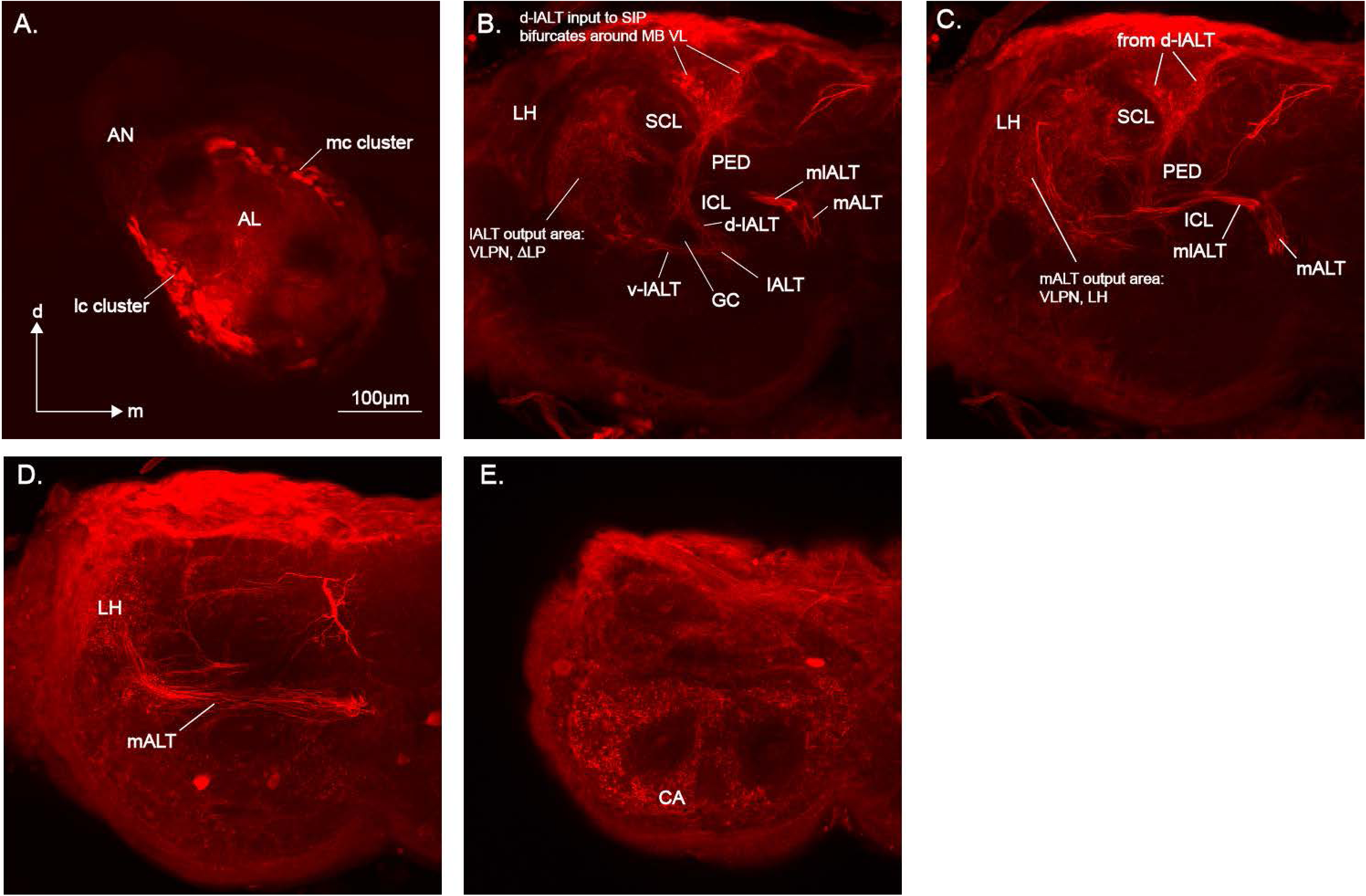
Mass staining of *H. virescens* male antennal lobe targeting lateral cell cluster neurons with dextran tetramethylrhodamine. A. Confocal images of the antennal lobe (AL) show that lateral cell cluster (lc) soma were primarily stained. Although some medial cell cluster (mc) soma were also stained, the patterns of projection to the protocerebrum were similar in all preparations. B. In this image projection neurons forming the diffuse lALT bifurcate around the Great Commissure (GC). The ventral aspect of the lALT pathway (v-lALT) proceeds to the lateral protocerebrum with terminals defining the Δ-lateral protocerebral area (ΔLP equivalent to the ΔILPC described in Seki et al. 2005; Namiki et al. 2014) located anteromedial of the Lateral Horn (LH). The ΔLP consists of parts of the ventrolateral protocerebral neuropils (VLPN: posterior ventral lateral protocerebrum PVLP, posterior lateral protocerebrum, PLP) and superior neuropils (superior lateral protocerebrum, SLP and superior clamp, SCL). The dorsal aspect of the lALT pathway (d-lALT) projects toward the dorsal surface of the protocerebrum along the superior intermediate protocerebrum (SIP) passing medial of the SCL and lateral of the pedunculus (PED) of the mushroom body before bifurcating in the dorsal SIP surrounding the vertical lobe of the mushroom body (VL). C. A number of neurons followed the mlALT passing above the inferior clamp (ICL) and then dipping below the PED before terminating both in an area that overlapped with lALT PNs in the ΔLP and also within the lateral horn (LH). D. PNs in the mALT continue toward the posterior of the protocerebrum before turning toward the lateral margin. In this image extensive terminal arborizations in LH can be seen. E. mALT PNs also arborize extensively in the calyces (CA) of the mushroom bodies. AN, antennal nerve, d dorsal, m medial. Scale bar = 100µm (all images).

**Figure 2.**
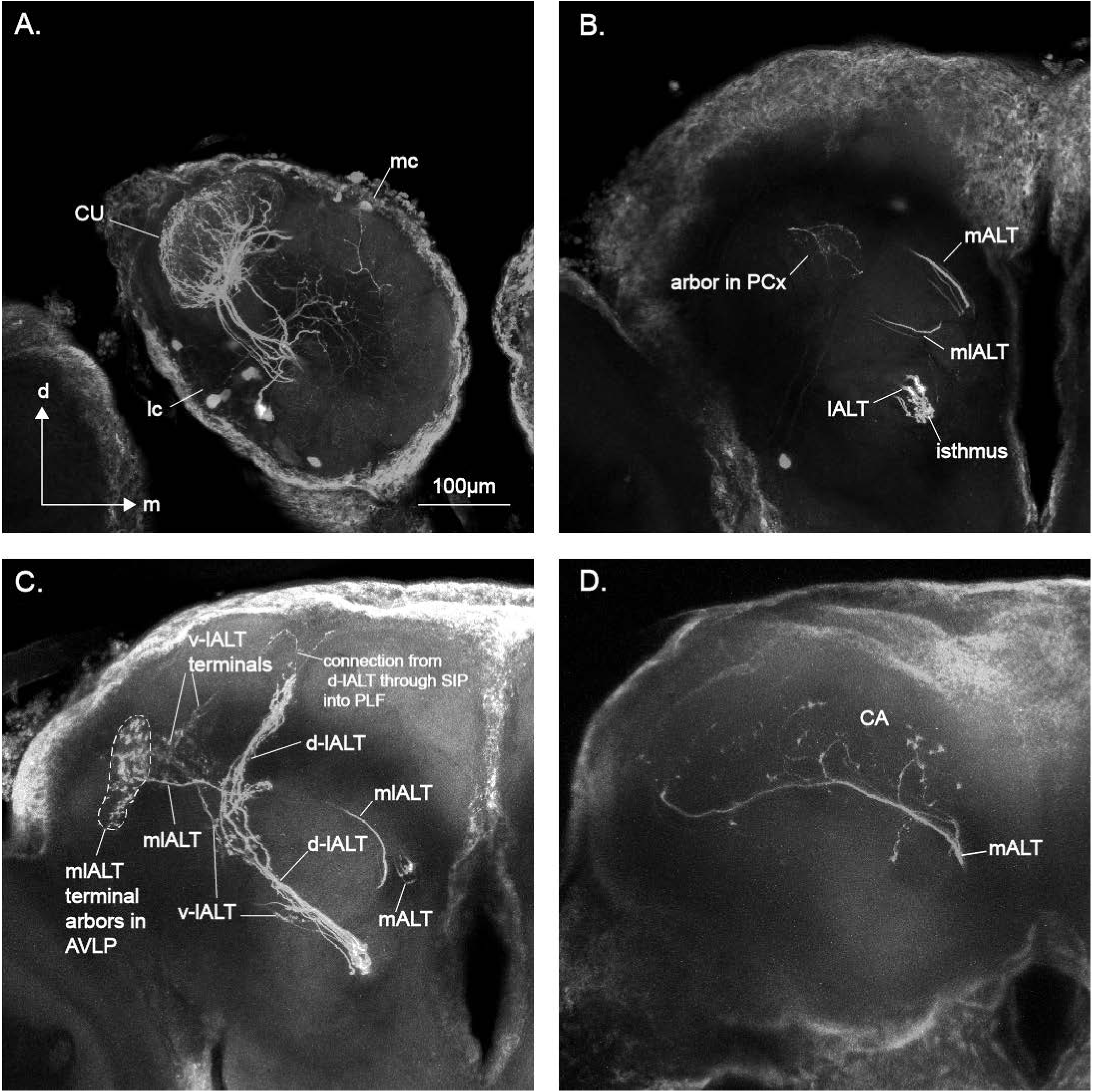
Indiscriminate staining of several *H. virescens* male antennal lobe projection neurons with neurobiotin revealed additional details of olfactory output pathways from *lc*PNs associated specifically with MGC glomeruli in the antennal lobe. A. Several lateral cluster (*lc*) somata were stained that gave rise to projection neurons. These PNs ramified in the MGC area particularly the cumulus (CU). Some medial cluster (*mc*) MGC PNs were also stained in this preparation including neurons associated with the cumulus and VM glomeruli. B. The three primary antennal lobe tracts can be seen clearly in this image. Axons that will form the mALT and mlALT are both heading toward the midline. Fibers constituting the lALT have reached the most medial aspect of their trajectory and some appear to make synaptic terminals in an area known as the isthmus before turning laterally. *lc*PNs clearly contributed to the mlALT and lALT tracts. C. The lALT diverged around the great commissure giving rise to a strongly stained group of fibers projecting dorsally (d-lALT) in the superior intermediate protocerebrum (SIP) alongside the vertical lobe of the mushroom body (VL) eventually splitting around the dorsal-most aspect of VL. Some of these fibers then appear to turn laterally and ventrally alongside the posterior lateral fascicle (PLF) and back toward the ΔLP area served by v-lALT PNs. The other, smaller group of lALT fibers in this preparation followed a pathway ventral of the great commissure (v-lALT) and proceeded toward the lateral protocerebrum before bifurcating into two main branches that define the ΔLP (=ΔILPC). One branch largely terminated in the area of the posterior ventrolateral protocerebrum (PVLP) but with some dendrites extending posteriorly into superior neuropils. Both v-lALT branches were anterior and medial of the lateral horn (LH). The dorsal and ventral lALT pathways thus intermingled along PLF. In this preparation, axons in the mlALT pathway turned toward the lateral protocerebrum and projected directly into an area consistent with the position of anterior ventrolateral protocerebrum (AVLP), just posterior of the antennal lobe and anteromedial of LH. Some dendrites appeared to spread posteriorly into PVLP overlapping with v-lALT *lc*PNs. D. mALT axons associated with MGC *mc*PNs proceeded caudally arborizing in the calyces of the mushroom body (CA) then turning posteriorly and dorsally before sending final branches into an area just posterior to the primary mlALT *lc*PN terminals in AVLP where they overlapped v-lALT terminals.

### Lateral Antennal Lobe Tract (lALT)

Staining in the lALT pathway was prominent but somewhat diffuse with neurons projecting to a variety of protocerebral targets (Fig. 1). The lALT turned sharply toward the lateral margin shortly after leaving the AL and bifurcated around the great commissure (GC). Axons following the dorsal route of the lALT (d-lALT) gave rise to a pathway that passed lateral of the inferior clamp (ICL) and pedunculus (PED) then medial of the superior clamp (SCL) before entering the superior intermediate protocerebrum (SIP) and bifurcating dorsally around the vertical lobe (VL) of the mushroom body. The aspect of the lALT pathway passing ventral of the GC (v-lALT) continued laterally toward the margin of the protocerebrum, terminating in an area anterior and ventral of the LH. The target areas and distribution of this lateral projection defined the extent of the ΔLP consisting of superior neuropils (superior clamp – SCL, and superior lateral protocerebrum – SLP) and inferior ventrolateral neuropils (posterior ventrolateral protocerebrum – PVLP, and posterior lateral protocerebrum – PLP) (Ito et al., 2014). Determining the precise relationships between terminal arborizations and specific neuropils in mass stains was challenging because of the indistinct boundaries between the separate neuropils that constitute the ΔLP and the ventrolateral protocerebral neuropils (VLPN) in general.

Neurobiotin staining (Fig. 2) revealed that some fibers departing through the lALT formed synaptic contacts as they exited the AL in an area identified as the isthmus in *M. sexta* (Homberg et al., 1988). Staining of l-*lc*PNs associated with the MGC cumulus was particularly strong in this preparation and permitted greater definition of pheromone-responsive *lc*PN targets. In this specimen, many fibers followed the d-lALT pathway to SIP. Fibers entering the lateral arm of the bifurcation around the VL portion of SIP appeared to continue dorsal of SCL and medial of the superior lateral protocerebrum (SLP) terminating alongside the posterior lateral fascicle (PLF) (Fig. 2C). Neurons following the v-lALT pathway with terminals in the posterior and dorsal portions of the ΔLP region also appeared to extend fibers along the PLF where they comingled with fibers from d-lALT neurons.

### Mediolateral-Antennal Lobe Tract (mlALT)

The mlALT was briefly confluent with the mALT as these tracts exited the AL, but soon diverged and projected toward the lateral margin of the PC (Figs. 1B, 2B,C). The common AL exit point for these two tracts was distinct from the lALT (Fig. 2B). Staining of the mlALT tended to be less prominent than the lALT, a finding consistent with a previous study estimating fewer axonal fibers in this pathway (Homberg et al., 1988). In dextran TMR mass-stained preparations, mlALT fibers passed dorsally of the ICL before dipping under the PED of the mushroom body (Figs. 1B,C). The tract continued dorsally and caudally to the anterior region of the lateral horn (LH) where terminal arborizations could be seen to comingle with projections from mALT PNs. mlALT *lc*PNs associated with the MGC and stained with neurobiotin revealed a distinct arborization in an area just posterior of the AL, anteromedial of LH, likely part of the anterior ventrolateral protocerebrum (AVLP) (Fig. 2C). In this preparation, fibers could also be seen extending more posteriorly and comingling with arbors from v-lALT MGC *lc*PNs in the ΔLP region (Fig. 2C).

### Medial Antennal Lobe Tract (mALT)

Staining of mALT fibers was also prominent in the mass-stain and neurobiotin preparations (Figs. 1D, 2B,D). Although *lc*PNs were the intended targets for mass staining, in almost all instances a few neurons with cell bodies in the medial cluster were also stained. Thus, the contribution of *lc*PNs to the mALT was not unequivocal. Nonetheless, the mALT certainly constitutes a well-characterized pathway that is followed by axons primarily derived from uniglomerular m-*mc*PNs. The mALT departs the AL close to the midline of the protocerebrum before diverging from the mlALT shortly thereafter and continues posteriorly before turning laterally and giving rise to dense arborizations in the calyces of the mushroom body (CA) (Figs. 1E, 2D) situated at the posterior margin of the protocerebrum. mALT fibers then turn anterodorsally with many terminating in LH (Fig. 1D). mALT *mc*PNs associated with MGC compartments (Fig. 2D) targeted an area medial and anterior of LH where they appeared to converge with v-lALT and mlALT *lc*PNs.

### *lc*PN physiology

Intracellular recordings from *lc*PNs were obtained in 9 preparations, 6 *H. virescens* and 3 *H. subflexa* (Figs. 3 and 4). Physiologies of 12 neurons were included in the *lc*PN physiology group on the basis of at least the presence of a cell body stain in the lateral cluster. Analysis of recordings revealed a diversity of physiological response profiles that were assigned to one of 2 broad categories: 1. Neurons that responded similarly to a single odorant and blends containing the same odorant (Fig. 3); 2. Neurons in which the responses to blends appeared different from single odorant responses (Fig. 4). For those neurons that responded to presentation of Z11-16:Ald alone, a comparison was made between responses to Z11-16:Ald and a pheromone mixture that included an equal amount of Z11-16:Ald. This analysis of 9 neurons (excluding the neuron that responded primarily to Z9-14:Ald, Fig. 3A, cell #1, and the two blend only neurons, Fig. 4C) revealed that, as a group, these cells responded to blends with significantly shorter response latencies and higher instantaneous spike frequencies (Fig. 5).

**Figure 3.**
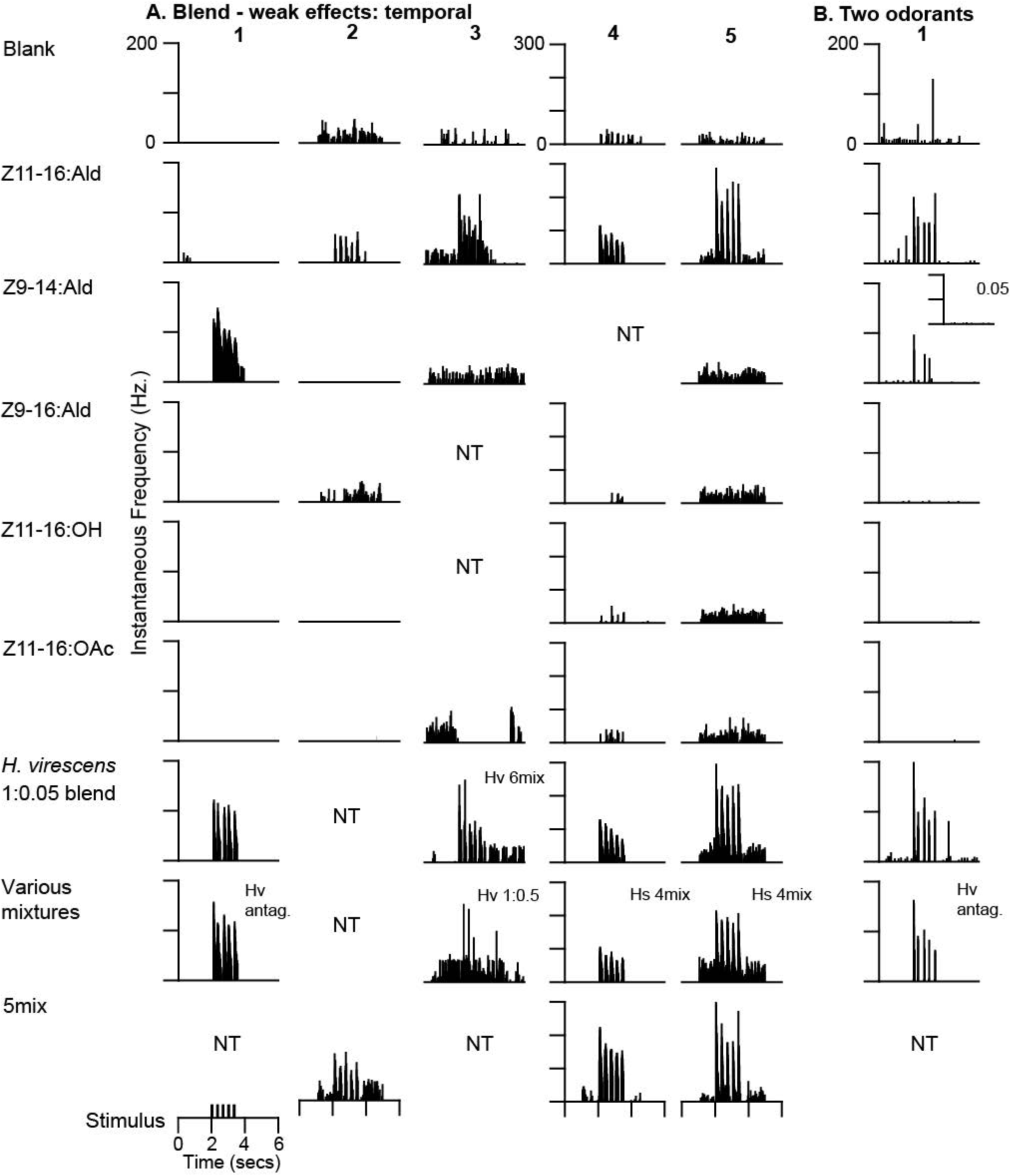
Moth *lc*PNs exhibited a diverse array of neurophysiological profiles as revealed by instantaneous frequency plots. Many *lc*PNs responded to presentation of a single odorant (Z11-16:Ald). A. Neurons were activated by presentation of brief pulses of Z9-14:Ald (cell #1) or Z11-16:Ald (cells #2-5). Responses to blends containing these odorants did not appear appreciably different (note in cell #1 the amount of Z9-14:Ald in each blend stimulus is considerably lower compared to the single odorant alone whereas in cell #s 2-5 the dosage of Z11-16:Ald is the same in single odorant and blend stimuli) B. A single *lc*PN responded to presentation of two odorants, more robustly to Z11-16:Ald than Z9-14:Ald (inset is response to low dosage, 0.05, of Z9-14:Ald used in *H. virescens* 1:0.05 blend). Blends containing both of these odorants did not appear to enhance or reduce the response compared to either odorant presented singly. NT, not tested. See Materials and Methods for notes on different mixtures used.

**Figure 4.**
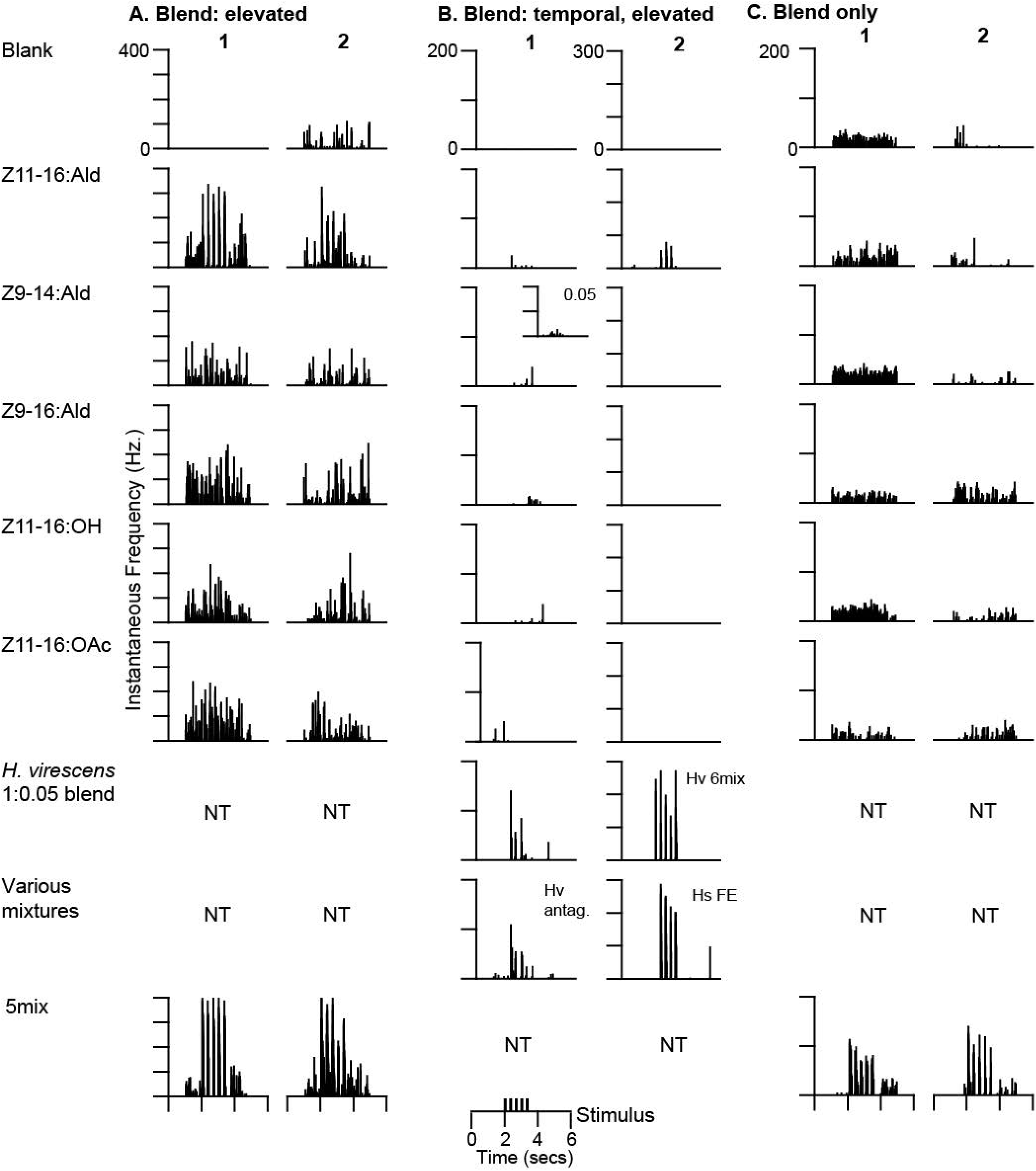
A. Two *lc*PNs responded to Z11-16:Ald but exhibited responses to blends in which the frequency of spiking increased. B. Recordings from two *lc*PNs revealed responses to a single odorant, Z11-16:Ald, but presentation of blend stimuli resulted in either higher frequencies of action potentials and/or more accurate following of the pulsatile stimulus. C. Two *lc*PNs either lacked or showed minimal responses to delivery of individual odorants but were clearly activated by pulses containing blends. NT, not tested.

**Figure 5.**
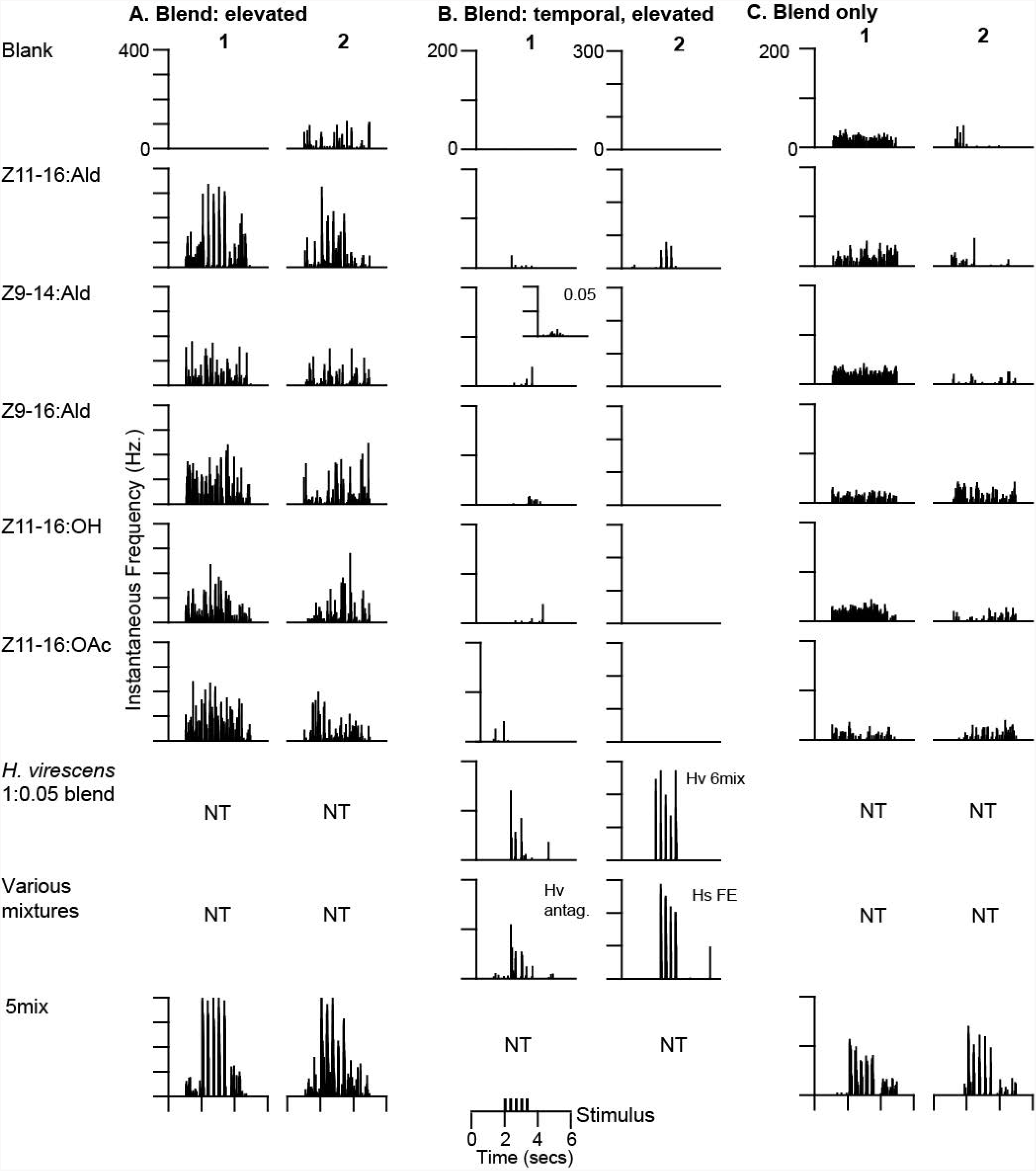
Analysis of *lc*PN physiological responses. A. Latency to onset of response was significantly reduced when *lc*PNs were challenged with a blend mixture versus Z11-16:Ald alone according to a matched pairs t-test (*P*=0.046). B. Peak instantaneous frequencies in response to a blend mixture containing Z11-16:Ald were significantly greater compared to responses to Z11-16:Ald alone (*P*=0.006).

### Single Odorant *lc*PNs

All neurons in this category responded to presentation with the major pheromone component, Z11-16:Ald, with the exception of one PN that exhibited a response to Z9-14:Ald. In the first response profile group of 5 PNs (Fig. 3A) excitatory activity elicited by blends appeared similar to responses elicited by a single odorant with perhaps some slight indication of elevated firing rates or improved pulse resolution. In a further category a single neuron was characterized as responding to both Z11-16:Ald and, more weakly, to Z9-14:Ald (Fig. 3B), with no obvious differences as a result of blend stimulation.

### Blend Response *lc*PNs

Two PNs (Fig. 4A) exhibited responses to Z11-16:Ald alone and blends containing this odorant were similar with respect to temporal resolution of odor pulses but responses to each blend stimulus pulse resulted in an elevated firing frequency compared to Z11-16:Ald alone. Two PNs exhibited responses that were enhanced (elevated excitation and/or temporal resolution of odor pulses) (Fig. 4B). Two additional PNs were identified in which responses to single odorants were not evident but presentation of one or more blends resulted in clear responses to each stimulus pulse (Fig. 4C).

### *lc*PN morphology

#### i. Glomerular associations

Detailed evaluation of *lc*PN morphology was possible in 8 of the 9 preparations in which physiological profiles were acquired (Figs. 6-11). A further 2 preparations in which no physiological records were obtained were included in the analysis of morphological features of *lc*PNs (Figs. 11A-C, 12). In 6 instances *lc*PNs projected from the AL via the lALT while in the remaining two preparations the PN axon followed the mlALT (Fig. 11). In all cases in which paired physiological and morphological information was obtained, dendritic arbors were observed in MGC glomeruli and always included the cumulus and DM. Preparations in which a single l-*lc*PN was stained revealed multiglomerular dendritic arbors present in both the cumulus and DM glomeruli (Figs. 6A-D; 8A-D). In several instances, the exact glomerular associations of individual *lc*PNs was unclear due to the staining of more than one neuron in the same preparation (Figs. 6 E,F, 7A-D, 9A). Nonetheless, in general, dendritic arbors of *lc*PNs were distinctly different from those previously reported for m-*mc*PNs. Uniglomerular m-*mc*PNs tend to exhibit dense dendritic arbors that define the boundaries of the glomerulus that they are associated with. In contrast, *lc*PNs tended to exhibit a sparser dendritic arborization that was most frequently characterized by the presence of several large branches around the perimeter of the glomerulus subsequently giving rise to smaller dendrites that penetrated the glomerulus interior (e.g. Figs. 6A,C, 8A, 10). This anatomy suggested the overall appearance of a broad basket around individual MGC glomeruli with a net of sparse smaller processes inside the glomeruli.

**Figure 6.**
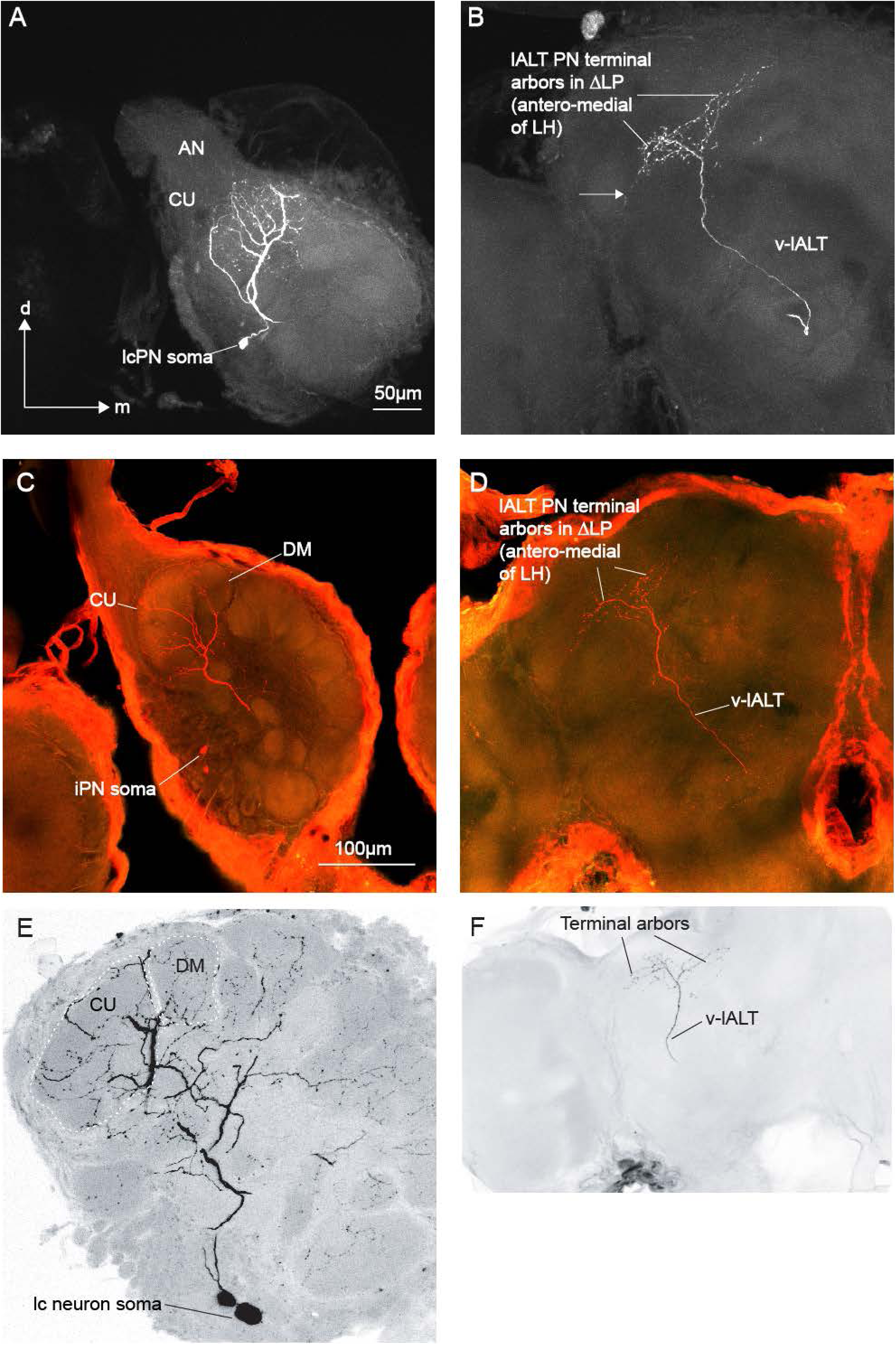
Three lateral cell cluster PNs with projections through the ventral pathway of lALT (v-lALT) exhibited similar morphological properties. A. In this preparation a single LY-stained *lc*PN exhibited dendritic arbors in both the cumulus (CU) and dorsomedial (DM) glomeruli of the MGC (dendritic arbors in the DM glomerulus were confirmed in sectioned material, images not shown). The basket-like dendritic arborization in the cumulus (CU) is most obvious in this projection of multiple images as compared to the dense dendritic arbor that characterizes typical uniglomerular mALT *mc*PNs. This *lc*PN responded robustly to stimulation with Z9-14:Ald and showed some improvement in following pulsatile stimuli in the presence of blends (Fig. 3A, cell #1). This was the only *lc*PN identified in the current study exhibiting a response only to Z9-14:Ald when presented as a single odorant (all others responding to Z11-16:Ald). B. The PN axon followed the v-lALT and made no apparent synaptic contact in the isthmus as it departed the AL before bifurcating in an area of the ventrolateral protocerebral neuropils (VLPN) anteromedial of the lateral horn with one branch invading the posterior ventrolateral protocerebrum (PVLP) (note also the small projection into the anterior ventrolateral protocerebrum, AVLP, arrow) with a second branch projecting more posterio-dorsally close to the posterior lateral fascicle (PLF) and proximal to superior clamp (SC) and superior lateral protocerebrum (SLP). This bifurcated pattern into two main branches that defines the ΔLP region (equivalent to the ΔILPC region described by Seki et al. 2005, and Namiki et al. 2014). Note a small amount of staining before the VLPN terminal arbors indicating a small projection into an area populated by d-lALT axons. C. A second *lc*PN stained with neurobiotin exhibited dendritic arbors in CU and DM. D. The axon followed the v-lALT and exhibited similar morphological characteristics as the *lc*PN featured in A, B. This neuron responded to Z11-16:Ald with no obvious effects of blends or mixtures (physiology: Fig. 3A, cell #5). E. In this *H. subflexa* male antennal lobe two *lc* neurons were stained with Lucifer yellow. The *lc*PN clearly had a dendritic arborization associated with the cumulus (CU). The other stained neuron was a local interneuron (iLN) that ramified throughout many of the AL glomeruli. The staining of two neurons made it difficult to determine if the *lc*PN had arbors in glomeruli other than the CU. F. The *lc*PN followed the v-lALT and exhibited terminal bifurcated arbors in the ΔLP similar to the other two preparations documented in this figure. This PN exhibited only weak responses to stimulation with Z11-16:Ald. Responses were greatly improved in terms of maximum frequencies and ability to follow delivery of all pulses in the presence of blends (physiology: Fig. 4B, cell #2).

**Figure 7.**
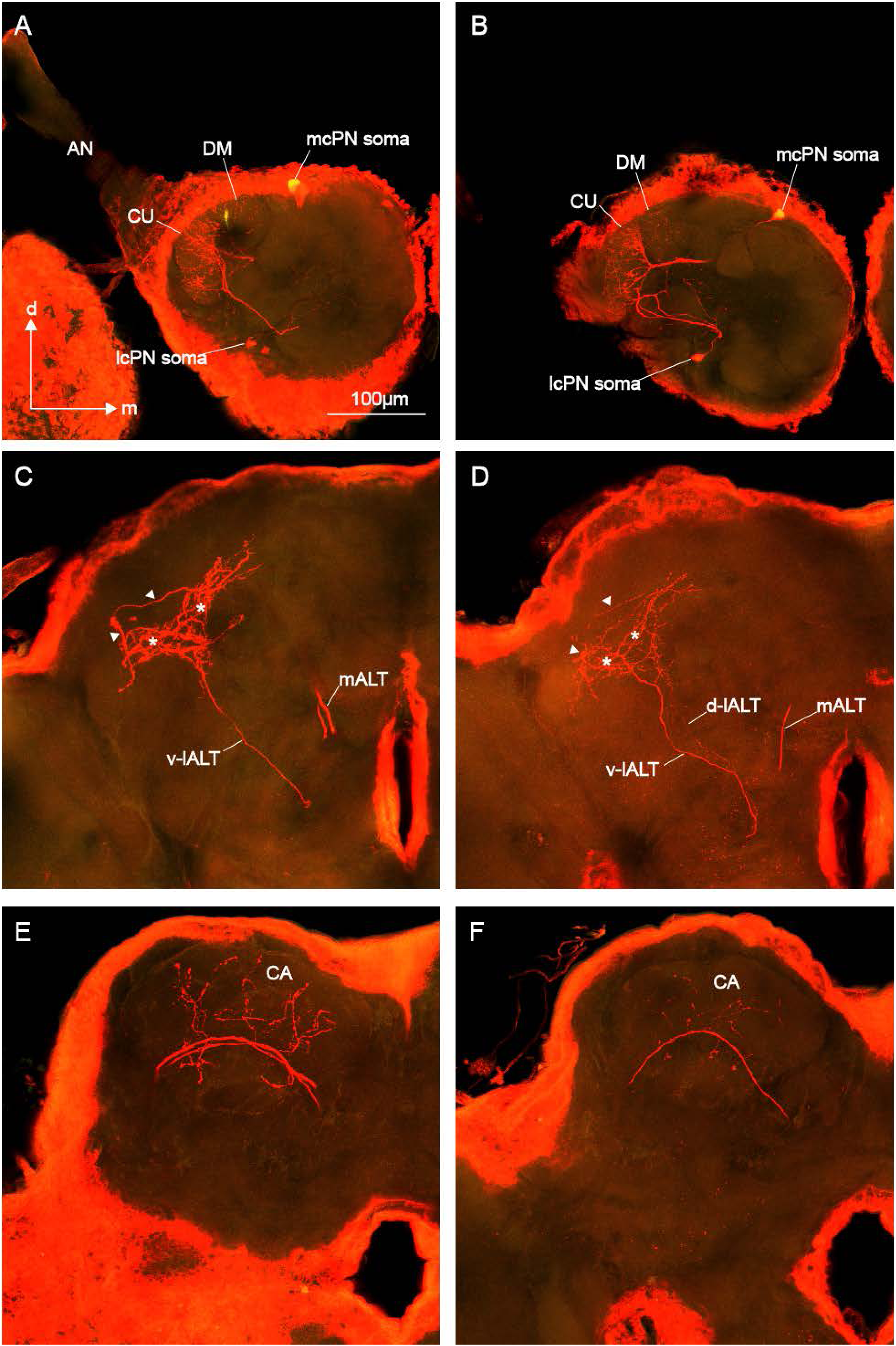
In two *H. subflexa* male preparations *lc*PNs and one or more mALT *mc*PNs with dendritic arborizations associated with the MGC were stained with neurobiotin. A, B. Both *lc*PNs appeared to have dendritic arborizations associated with the cumulus (CU) and dorsomedial (DM) MGC glomeruli. The m-*mc*PNs appeared to have strong uniglomerular dendritic arborizations associated with CU. C, D. The *lc*PNs exhibited a similar pattern of output arborization as neurons described in Fig. 6 i.e. axons followed the v-lALT pathway with a birfurcated arbor anteriomedial of the LH defining the ΔLP region. Note that in both cases the m-*mc*PN lateral terminals also exhibited two main branches (arrowheads) extending into the same protocerebral ΔLP areas as the branches of the v-lALT *lc*PNs (*). E, F. m-*mc*PNs neurons exited the AL through the mALT and proceeded posteriorly to the mushroom body calyces (CA) where they exhibited extensive output arborizations before turning anteriorly. In one preparation (images A, C, E), the physiology of a single neuron was captured (Fig. 3A, cell #4). This neuron responded robustly to presentation of Z11-16:Ald with similar responses to various mixtures containing this odorant.

**Figure 8.**
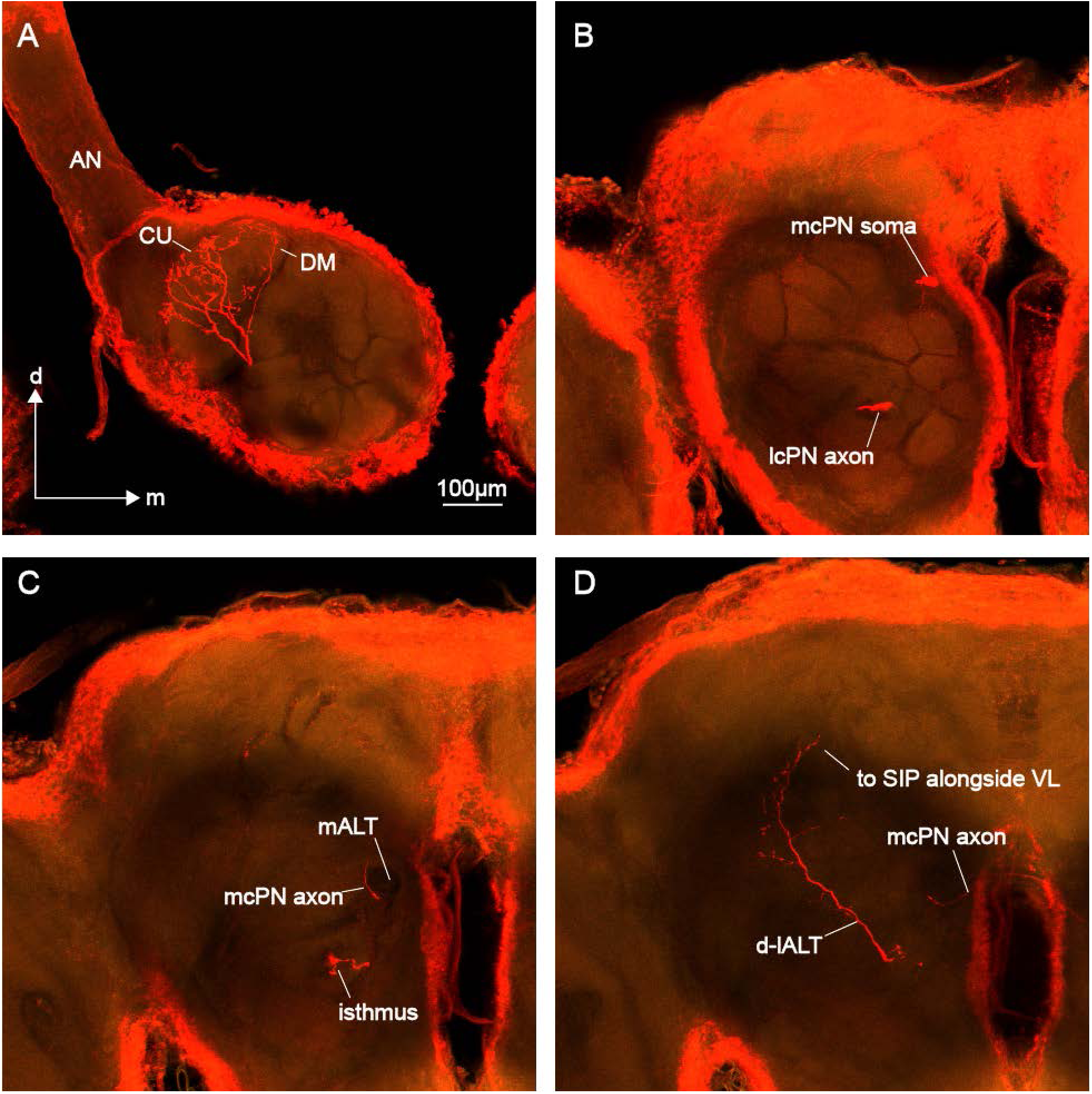
l-*lc*PNs that follow the d-lALT exhibited distinct morphological features compared to v-lALT PNs. A. This particular l-*lc*PN arborized in MGC glomeruli cumulus (CU) and DM. Again note that the dendritic branching pattern is relatively sparse in comparison with canonical uniglomerular m-*mc*PNs. B. A second neuron (*mc*PN) was stained but appeared to have no dendritic arbor in the ipsilateral AL. C. As the l-*lc*PN axon departed the antennal lobe clear synaptic swellings were observed in the isthmus area. The *mc*PN did not project out of the AL through any of the three large ipsilateral ALTs. Instead its axon skirted the mALT before crossing the midline just above the subesophageal ganglion. D. The l-*lc*PN axon proceeded along the dorsal pathway of the lALT (d-lALT) before turning dorsally, lateral of the pedunculus into the superior intermediate protocerebrum (SIP) alongside the vertical lobe of the mushroom body. Note that the initial length of the axon beyond the isthmus does not appear to have any en passant synaptic swellings which then appear distally. The physiology of this blend elevated neuron is shown in Fig. 4A (cell #2).

**Figure 9.**
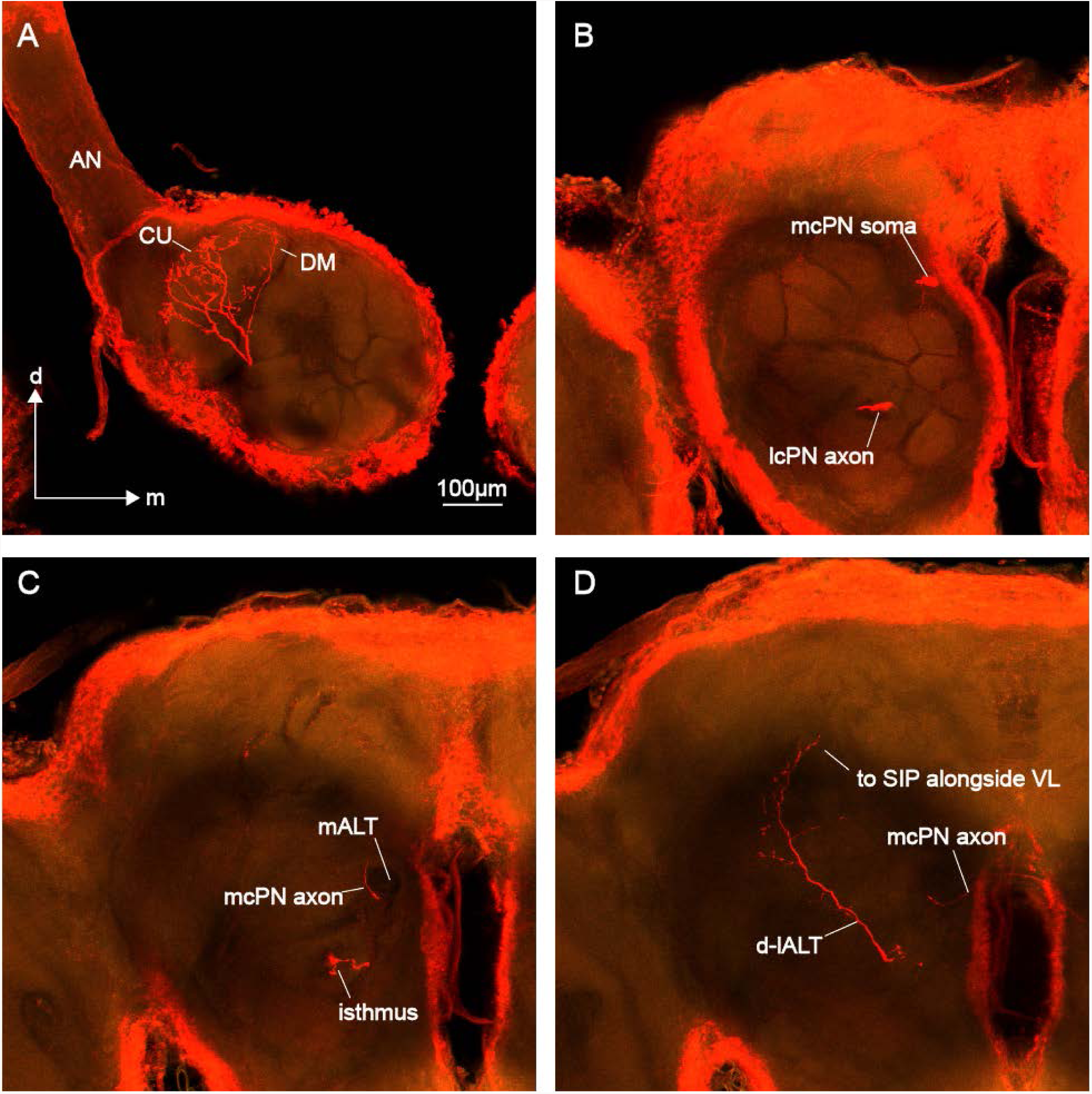
Diverse morphologies of l-*lc*PNs. A. In this preparation several l-*lc*PNs were stained with neurobiotin as well as three m-*mc*PNs. Stained *lc*PNs arborized extensively within the cumulus (CU) of the MGC with at least one neuron also arborizing in DM. One iLN was also stained. One of the m-*mc*PNs was clearly associated with the MGC VM glomerulus (axon could be traced to the mushroom body calyx and ΔLP where it overlapped with l-*lc*PN terminals) while another arborized within a medially-located ordinary glomerulus (OG, axon not fully stained). B. The *lc*PNs projected from the AL via the lALT with some forming synaptic contacts in the isthmus. The stain in the dorsal protocerebral area (arrow) originates from contralateral lcPN shown in (E). C. lALT tract shortly after turning toward the lateral protocerebral margin. *lc*PNs followed either the v-lALT or d-lALT pathways. v-lALT neurons bifurcated around GC and terminated in the ΔLP region. d-lALT neurons turned toward the dorsal margin along the vertical lobe of the mushroom body. Lateral of GC the axon of one v-lALT PN turned toward the midline and gave rise to a dorsal branch that partially encircle the pedunculus (PED). Some of these fibers followed PED back to its origin in the mushroom body (not shown). The main axon continued toward the midline. D. Arborizations in the CA of one m-*mc*PN, staining in the lateral PC from one or more of these m-*mc*PNs was restricted to the LH (not visible in this image) suggesting that this *mc*PN was associated with stain in an ordinary glomerulus in the antennal lobe. E. Composite of two confocal images one from each side of the protocerebrum showing the ipsilateral *lc*PN projections through v-lALT and d-lALT on the right of the protocerebrum and the contralateral projection of one v-lALT *lc*PN. The axon projects across the midline to the opposite protocerebral hemisphere. Note the symmetrical branches that arise just before either side of the midline (arrows, also arrowed in B on the ipsilateral side) and curve back laterally through the superior protocerebral neuropils and into the region invaded by ipsilateral d-lALT *lc*PNs. Note the symmetrical morphology and the blebby synaptic processes on the contralateral side that appear to run along the contralateral d-lALT extending both anteriorly to the isthmus as well as dorsally toward the contralateral pedunculus. Staining in an area anteromedial of the LH in the contralateral hemisphere originates with an MGC m-*mc*PN from the ipsilateral antennal lobe. The physiology from one blend elevated neuron in this preparation was recorded (Fig. 4A, cell #1).

**Figure 10.**
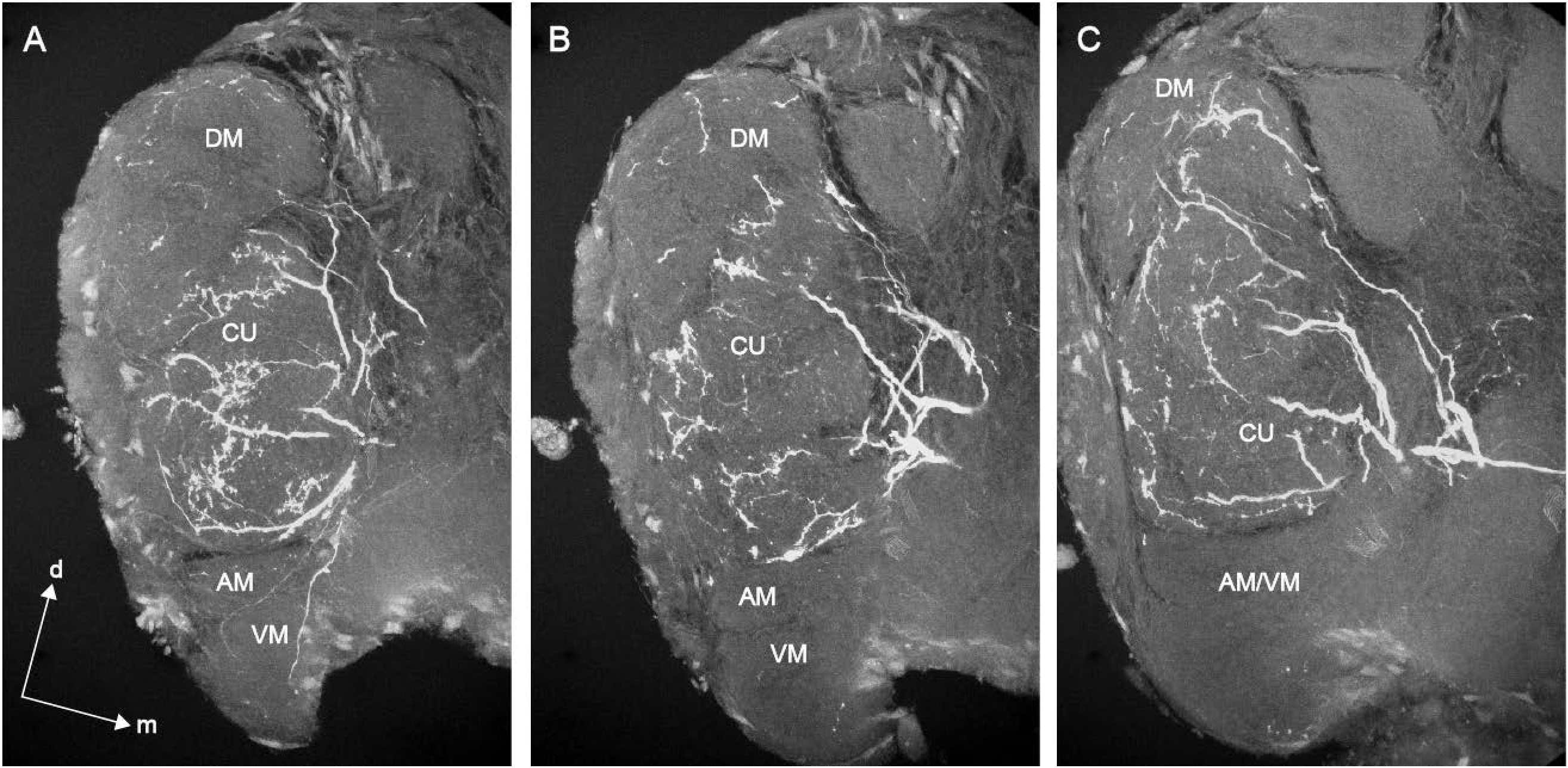
Confocal images of a sectioned antennal lobe in an *H. virescens* male captured with a 40X objective reveal dendritic arborizations of *lc*PNs stained with LY in detail. A. Dendritic architecture appears sparse with several processes positioned around the glomerular edge with a few large arbors located within the glomerular neuropil. In this image from more anterior confocal sections, arborizations are prevalent in the cumulus with a few located in the dorsomedial glomerulus (DM) of the MGC. Processes skirted the borders of the anteromedial (AM) and ventromedial (VM) MGC glomeruli (arrowheads) but did not appear to penetrate the glomerular neuropil. B. More processes become visible in DM. Many of the processes in the cumulus are located at the glomerular periphery. No arborizations are visible in AM/VM. C. Arborizations are present in the cumulus and DM. Three *lc*PN soma were observed in this preparation although only one PN appeared to be fully stained. Three similar physiological records were acquired but the responses of only one neuron are included (Fig. 3A, cell #3). d, dorsal, m, medial.

#### ii. *lc*PNs projecting through lALT

*lc*PNs following the v-lALT pathway made no synaptic contact in the isthmus (Figs. 6B,D, 7C,D). Their projection into the lateral protocerebrum terminated in a bifurcated arborization defining the *Δ*LP, one anterior branch likely in the posterior part of AVLP and PVLP anteromedial of LH and a second branch extending posteriorly and dorsally with arbors likely in PLP, SCL and along PLF, medial of LH. These *lc*PNs also frequently had some staining in the vicinity of the dorsally projecting pathway of the d-lALT neurons (Figs. 6B,D) and did not penetrate LH. In those preparations in which an MGC m-*mc*PN was also stained, it was evident that the terminal arborizations of the l-*lc*PNs and m-*mc*PNs overlapped (Fig. 7C,D) in both anterior and posterior regions of the *Δ*LP. Thus, MGC m-*mc*PNs also did not terminate in the LH. In contrast, d-lALT *lc*PNs exhibited a clear area of synaptic contact in the isthmus as their axonal fibers departed the AL (Figs. 8C, 9B). Furthermore, these *lc*PNs had many swellings along their axon indicative of synaptic contacts. These axons turned dorsally and appeared to run alongside VL within SIP alongside the anterior optic tubercule (AOTU) (Fig. 8D). This part of the SIP was termed the *column* by Ian et al. (2016b) although this is not a recognized structure in the consensus insect brain (Ito et al. 2014).

#### iii. *lc*PNs projecting through the mlALT

Two *lc*PNs had projections through the mlALT to the *Δ*LP (Fig. 11). Both neurons exhibited similar patterns of arborization within the AL – thin, weakly stained processes that invaded both the cumulus and DM glomeruli of the MGC (Figs. 11A,B). Axons from these PNs exited the AL alongside the mALT but then turned laterally. Terminal arborizations consisted of two main branches – one branch reaching into an area just posterior of the AL, likely AVLP, with the other branch projecting posteriorly and dorsally into an area that is also targeted by v-lALT *lc*PNs (Figs. 11C,D).

**Figure 11.**
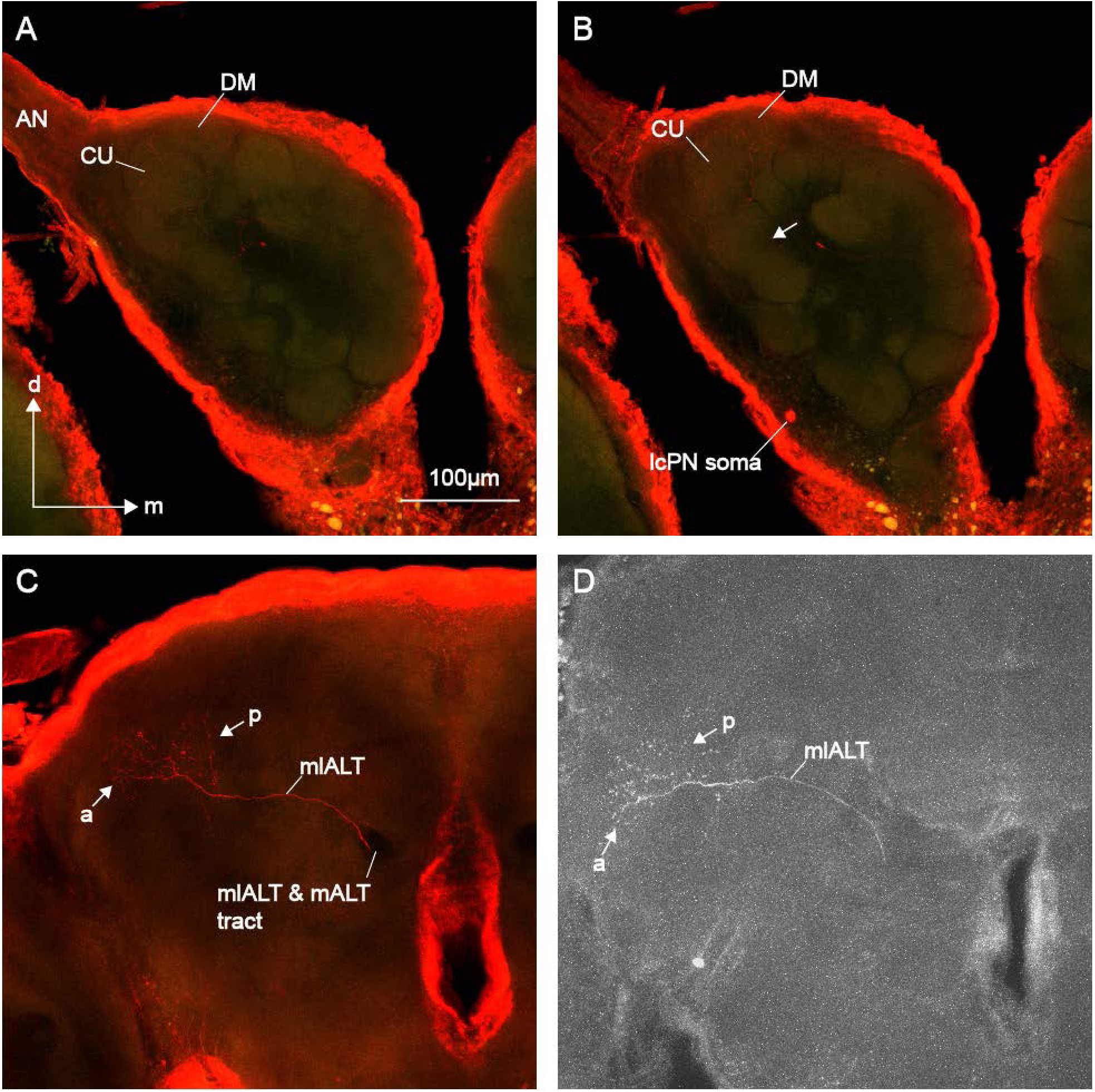
*lc*PNs projecting via the mlALT to the lateral protocerebrum were less commonly encountered, likely indicative of the smaller number of neurons associated with this tract (Homberg et al. 1988). In these two preparations, arborizations within the AL were only barely visible. A, B. In this preparation, an *lc*PN and iLN were both weakly stained with neurobiotin. Arborizations of the *lc*PN were associated with the cumulus (CU) and DM. Neurite of the *lc*PN is indicated by an arrow (brighter neurite originates from the iLN). C. The mlALT and mALT are congruent as they leave the AL but the mlALT soon deviates toward the lateral PC. This neuron appeared to have a bifurcated terminal arborization in the VLPN area, one branch anterior (a) at least partially in the anterior ventrolateral protocerebrum (AVLP) and the other posterior (p) overlapping with v-lALT *lc*PN terminals in ΔLP. D. Protocerebral arborizations of a second mlALT *lc*PN stained with LY exhibited very similar characteristics to the neuron depicted in A-C, weak staining in the AL (CU and DM – not shown), with stronger protocerebral staining. Physiology for this neuron in Fig. 3B, cell #1. This neuron responded to two odorants, Z11-16:Ald and Z9-14:Ald with little difference noted to stimulation with blends. d, dorsal, m, medial.

#### iv. *lc*PNs projecting through mALT

In one instance an *lc*PN associated with the posterior complex (PCx – Lee et al 2006a,b) an area just posterior of the MGC displayed a dense tuft arborization reminiscent of an m-*mc*PN (Fig. 12B). Interestingly, the axon of this PN exited the AL through the mALT and exhibited widespread terminal arborizations in the CA and LH that broadly overlapped with an m-*mc*PN (with dendritic arbors associated with an ordinary glomerulus) that had been stained in the same preparation (Fig. 12C,D).

**Figure 12.**
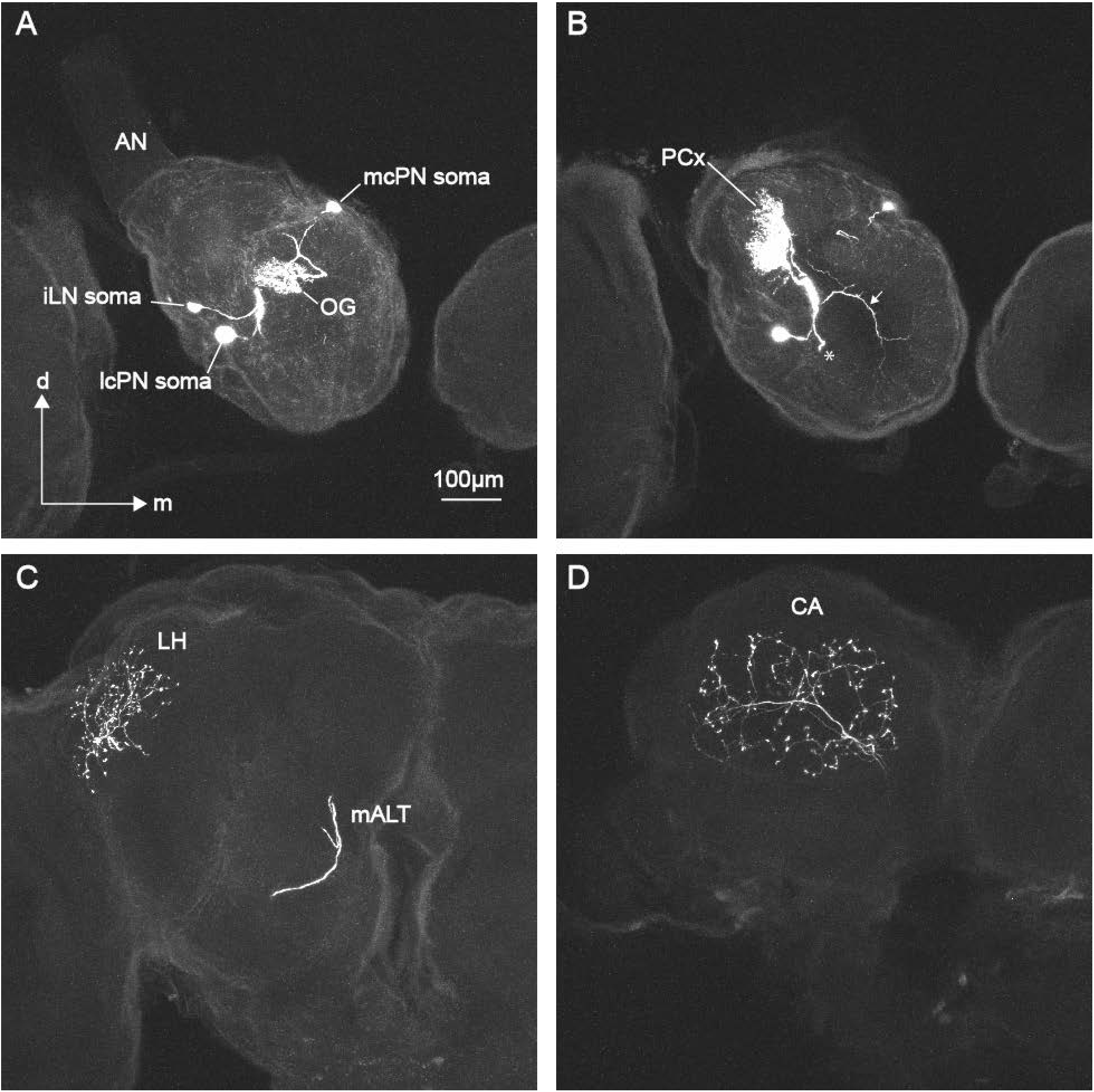
*lc*PNs also projected to higher brain centers through the mALT. A. Three neurons were stained with LY in this preparation (no physiology was obtained for these neurons). Two *lc* neurons and one *mc*PN were stained. One *lc*PN was strongly stained whereas the other lc soma was associated with a wide-field local interneuron (iLN). In this image, the *mc*PN soma and arborization (restricted to a single ordinary glomerulus, OG) can be seen. B. The dendritic arbor of the *lc*PN was restricted to the posterior portion of the AL, in two glomeruli that form part of the so-called posterior complex (PCx). Note the dense dendritic tuft reminiscent of those associated with canonical uniglomerular m-*mc*PNs. The axon of the *lc*PN is indicated by an * whereas the iLN primary neurite is indicated by an arrowhead. C. Both OG m-*mc*PN and PCx *lc*PN departed the AL through the mALT. In this image overlapping terminal arborizations can be seen in the lateral horn (LH). D. Both PNs also exhibited extensive overlapping ramifications in the calyces (CA) of the mushroom body.

#### V. Contralateral projections

In two preparations contralateral projections were noted, one from an *lc*PN and the other from an *mc*PN (Figs. 8D; 9E respectively). The morphology of the contralateral *lc*PN (Fig. 9E) is similar to ones previously reported in *A. segetum* (Wu et al. 1996) and *H. virescens* (Ian et al. 2016b). The axon of this PN followed the v-lALT ipsilaterally with no apparent synaptic contacts in the isthmus or along the ipsilateral process (Fig. 9E). After passing ventral of the GC, the axon turned back toward the midline, and appeared to send a process dorsally to make contact with the outer margin of the pedunculus. The main axon continued toward the midline dorsal of the central body and a thin process emerged from it just before the midline. This process projected back into the target area of d-lALT PNs in SIP and blebby processes indicated synaptic contacts in this area. This PN had a generally symmetrical appearance but the contralateral projection ran both ventrally to the isthmus and dorsally toward the pedunculus along the d-lALT and not the contralateral v-lALT. Swellings along these contralateral ramifications indicated the presence of synaptic connections in these areas (including the isthmus area) (Fig. 9E).

### *lc*PN combined morphology/physiology

In several preparations more than one neuron was stained and, as such, unambiguous associations between *lc*PN neuronal staining and physiological records was limited. Nonetheless, our data support the following conclusions: multiglomerular MGC *lc*PNs (Figs. 6A-D, Fig. 3A cell #s 1&5) projecting through the v-lALT included responses to single odorants (Z9-14:Ald and Z11-16:Ald respectively) in their repertoires with similar responses to stimulation with blends. Another v-lALT PN exhibited weak responses to Z11-16:Ald, but greatly elevated and temporally enhanced responses to pheromonal blends (Figs. 6E-F, Fig. 4B cell #2). These neurons all exhibited very similar bifurcated terminal branches defining the *Δ*LP region with an anterior branch likely in AVLP/PVLP and a posterior, dorsal branch projecting into PLP and dorsally into SLP and SCL alongside PLF. In all of these preparations there is evidence of some staining in the same area followed by dorsally projecting d-lALT PNs. Another MGC *lc*PN (Figs. 7 A,C,E; Fig. 3A cell #4) also projected along the v-lALT and exhibited a similar *Δ*LP axonal arborizations. These terminals appeared to overlap extensively with an MGC m-*mc*PN stained in the same preparation. The l-*lc*PN neuron responded to Z11-16:Ald and appeared to have a slightly elevated response to a pheromone mixture. A multiglomerular (cumulus and DM) *lc*PN (Figs. 8A-D) stained with neurobiotin exhibited an area of synaptic contact in the isthmus before projecting along d-lALT to SIP (physiology: Fig. 4A, cell #2). Another two PNs from this preparation responded exclusively to blends (Fig. 4C) (one neuron was recorded from the right AL while the other was in the left AL, both with microelectrodes containing LY). *lc*PNs that responded to presentation only of blends tended to be MGC multiglomerular (primarily cumulus and DM), exhibited an area of putative synaptic contact in the isthmus as the axon exited the AL and then projected along the d-lALT before turning dorsally toward the SIP in the dorsal PC. Unlike v-lALT *lc*PNs these neurons also exhibited blebby synaptic processes along most of their axonal length. A paired recording/stain was obtained from a single ml-*lc*PN (protocerebral projection shown in Fig. 11D, physiology: Fig. 3B) with very weak multiglomerular staining in the MGC (cumulus and DM). This neuron responded to separate stimulation with two pheromonal odorants (Z11-16:Ald and Z9-14:Ald) and similarly to blends thereof. This ml-*lc*PN had similar AL morphological characteristics to the PN shown in Figs. 11A-C for which no physiology was recorded.

## Discussion

The existence in moths of a parallel pathway for olfactory information to the protocerebrum convergent with outputs of excitatory PNs is consistent with studies on olfactory processing in other insect taxa including fruit flies and honeybees (Galizia and Rössler 2010). As with other studies (Kanzaki et al., 2003; Rø et al. 2009, Ian et al. 2016), we confirm that in moths this parallel pathway originates with olfactory PNs having cell bodies located in the lateral cell cluster of the moth AL. This AL output travels along several different routes to higher brain centers. In addition to the varied morphological features of these neurons we were able to document a diversity of neurophysiological response profiles contrasting with the relatively predictable characteristics of their mostly uniglomerular m-*mc*PN counterparts. Dendritic arborizations of all ml-*lc*PNs and l-*lc*PNs reported here were associated with the MGC except for one m-*lc*PN with dendritic arborizations in PCx. While some neurons were activated by single odorants primarily the main pheromone component (Z11-16:Ald), in many cases responses were improved by stimulation with a behaviorally relevant pheromone blend. Based on limited data, Baker and Hansson (2016) proposed that the ml-and l-ALT pathways relayed ‘chromatic’ information (i.e. activity dependent on presence of a behaviorally active pheromone blend) to the lateral protocerebrum but suggested that these neurons exhibited poor temporal resolution of pusatile stimuli. The current study makes it clear that the response profiles of these neurons span from achromatic (similar responses to a single odorant and blends) to fully chromatic (only blend elicited activity). In addition, the temporal resolution of *lc*PNs as a group was measured by their ability to track pulsatile stimuli and was found to be significantly enhanced by presentation of blend stimuli compared to single pheromonal odorants alone.

Dendritic arborizations of all l-*lc*PNs and ml-*lc*PNs occurred with the MGC cumulus plus DM glomeruli. We were not able to resolve any single ml- or l-*lc*PNs restricted only to the cumulus although Vickers et al. (1998b) reported one such example in *H.virescens* (two overlapping v-lALT uniglomerular blend synergist *lc*PNs reported in Vickers et al. 1998 was mistakenly identified as following the mlALT). However, the wide variety of *lc*PN response profiles did not simply reflect the relative homogeneity of their arborization patterns. In contrast, the majority of m-*mc*PNs convey information related to activity in a single glomerular location to CA/LH and, as such, their responses to single odorants are often similar to those elicited by blends. However, there are also examples of multiglomerular blend sensitive m-*mc*PNs included amongst them (e.g. *H. virescens* blend-enhanced response shown in Fig. 12, Vickers et al. 1998)

It is clear from the current study that pheromone-responsive moth *lc*PNs have widespread ramifications in the protocerebrum. Moth MGC *lc*PNs exited the AL along all three main AL output tracts, although the majority reported here followed lALT with some in mlALT and only one example of a mALT PN. The lALT has previously been identified as a major output pathway for *lc*PNs (Homberg et al., 1988; Berg et al., 2007; Ian et al., 2016a,b). The current study confirms that pheromone-responsive *lc*PNs contribute to the lALT but also reveals that this tract bifurcates around the GC giving rise to ventral and dorsal pathways (Rø et al. 2007; Ian et al., 2016b). Physiologically these PNs responded to single odorants but were characterized as blend enhanced. The primary targets of moth MGC v-lALT *lc*PNs are in a pyramidal-shaped area anteromedial of LH, the ΔLP (previously identified as ΔILPC in *B. mori,* Kanzaki et al., 2003; Seki et al., 2005; Namiki et al. 2013). Results from *B. mori* indicate that MGC m-*mc*PNs have different patterns of arborization in CA according to the MGC glomerulus with which their dendritic arbors are associated (Kanzaki et al., 2003; Seki et al., 2005; Namiki et al., 2013; reviewed in Baker and Hansson, 2016) and project to different regions of ΔLP where they overlap with l-*lc*PNs. Our results concur that MGC m-*mc*PNs and l-*lc*PNs have overlapping projections in ΔLP but we were unable to resolve if uniglomerular m-*mc*PNs associated with different MGC glomeruli exhibited a separation in their terminal arborizations. However, results from a related heliothine moth, *Helicoverpa assulta* indicate that such a segregation is likely (Zhao et al., 2014).

Multiglomerular l-*lc*PNs followed the d-lALT with a dorsal axonal projection adjacent to the VL in SIP as also observed in mass stainings (Ian et al., 2016b). In addition, only d-lALT *lc*PNs formed areas of synaptic contacts within the isthmus as the lALT departed the AL suggesting that these neurons may either convey or receive information from mechanosensory pathways that passage through this structure (Homberg et al., 1988). These neurons also exhibited possible en passant synaptic swellings along their axons. Their response profiles were characterized as mostly blend dependent. Rø et al. (2007) reported a similar l-*lc*PN with sparse dendritic arborizations in several glomeruli from a female *H. virescens* indicating that the d-lALT pathway is not restricted to pheromone processing in males. Another such l-*lc*PN was reported by Ian et al. (2016a) although it was not clear if this neuron was stained in a female or male *H. virescens*.

Contralateral neurons of the type reported here (Fig. 9E) have only been noted occasionally in *Manduca sexta* (Homberg et al., 1988), *H. virescens* (Ian et al., 2016a) and another noctuid species, *Agrotis segetum* (Wu et al., 1996). Our results indicate that the contralateral neuron follows the v-lALT before sending a branch toward the margin of PED with the main axon turning toward the midline. Iwano et al. (2010) reported that the lALT skirts around the medial margin of the lateral accessory lobe (LAL) before turning around its posterior edge toward the lateral protocerebrum. Since v-lALT *lc*PNs did not exhibit en passant axonal synapses, this makes it unlikely that these contralateral neurons have synaptic contacts within LAL as Wu et el. (1996) concluded from their study. The contralateral *lc*PN reported here was also not entirely symmetrical in that presumptive terminal synaptic connections appeared within the contralateral d-lALT, as well as the isthmus and toward the peduncular region although the precise targets of these terminal connections were not clear.

Recent work in *Drosophila melanogaster* has highlighted the physiological, anatomical and potential behavioral significance of olfactory *lc*PNs associated with the fly AL ventral cell cluster (*vc*) (Liang et al., 2013; Parnas et al., 2013; Strutz et al., 2014; Wang et al., 2014). In flies the functional equivalent of neurons with cell bodies in the moth *lc* appear to be split into two separate cell body clusters, the *vc* housing inhibitory GABAergic PNs (iPNs) and a dorsolateral cluster (*dl*) containing inhibitory LNs (iLNs) (Tanaka et al. 2012). Immunohistochemical studies support the conclusion that all neurons with cell bodies in the moth *lc* are also GABAergic (*M. sexta*: Hoskins et al., 1986; *H. virescens*: Berg et al., 2009). Most fly iPNs exit the AL via the mlALT (ml-iPNs) and project to the LH and VLPN (Tanaka et al., 2012). A third cell cluster situated anteriodorsally (*ad*) of the AL in *D. melanogaster* harbors primarily uniglomerular excitatory, cholinergic PNs, the functional equivalents of moth m-*mc*PNs that similarly exit the AL via the mALT (m-ePNs). As in moths, the initial primary target of the fly m-ePNs is the CA before progressing to final terminal arborizations in LH. Recent studies into the functional role of fly ml-iPNs have reached a variety of conclusions about the role of these neurons in processing odorants either associated with food and/or conspecifics (Liang et al., 2013; Parnas et al., 2013, Wang et al., 2014, Strutz et al., 2014). However, a synthesis of these results into a consensus understanding of iPN connectivity and circuit function in the protocerebrum has yet to emerge.

Only two neurons were stained with a projection through the mlALT and mass staining confirmed that this pathway is comprised of fewer fibers than the lALT in moths (Homberg et al., 1988), the opposite of the relationship between these two tracts in *Drosophila*. A neurophysiological profile was obtained for one neuron and revealed responses to two single odorants (Z11-16:Ald and Z9-14:Ald) but did not appear to be blend enhanced. Both neurons appeared to receive input in the MGC but a lack of strong staining in the AL prevented an unambiguous determination of glomerular associations. In the moth brain the mlALT pathway appears to play a reduced role in central pheromone processing compared to the lALT.

We noted that dendritic glomerular aborizations of ml- and l-*lc*PNs in the moth AL were often sparser than the dense uniglomerular arborizations of canonical m-*mc*PNs. This morphology is consistent with observations from the fly (Strutz et al., 2014; Tanaka et al., 2012b) and may indicate a distinct synaptic connectivity. Morphological evidence also supports the conclusion that some *lc*PNs exit the AL via the mALT alongside m-*mc*PNs with which they have overlapping projections in CA and LH. Interestingly, one such m-*lc*PN stained during the course of the current study exhibited a dense uniglomerular dendritic tuft within a PCx glomerulus (Lee et al., 2006a,b). In female moths, Reisenman et al. (2004) reported a linalool-responding uniglomerular *lc*PN in *M. sexta* that directly targeted CA/LH while Løfaldi et al. (2010) described the morphologies of one uniglomerular and one multiglomerular m-*lc*PN from female *H. virescens* both with terminals in CA and LH. As such, stains of *lc*PNs associated with retrograde mass stains in CA reported by Ian et al. (2016b) may at least partially be due to these m-*lc*PNs.

Not all MGC glomeruli were innervated by moth ml-*lc*PNs and l-*lc*PNs. Multiglomerular *lc*PNs always invaded both the cumulus and DM and dendritic arbors were sometimes observed to skirt around AM and VM but there was no direct evidence for penetration of the glomerular neuropil in either *H. virescens* or *H. subflexa* males. In *H. subflexa*, the AM glomerulus receives inputs from OSNs tuned to Z11-16:OH – an essential component of the pheromone of this species (Lee et al., 2006b). Thus, combinations of activity within the cumulus and DM are important in the transmission of information, with likely inhibitory synaptic connectivity, to higher brain centers. Our results do not shed light on the exact synaptic partners of ml- and l-*lc*PNs in ΔLP. It is possible that pheromone sensitive m-*mc*PNs and *lc*PNs interact directly with each other in these areas but it seems more likely that they converge on a common target neuronal population to either influence or gate the passage of olfactory-driven activity to tertiary circuits. The area encompassed by ΔLP includes small superior-most parts of PVLP and PLP – both areas reported to contain optic glomeruli that receive visual input from the lobula plate of the optic lobe (Ito et al., 2014; Strausfeld and Okamura, 2007; Strausfeld et al. 2007). Baker and Hansson (2016) proposed that ΔLP could be an important neural substrate for multimodal integration of olfactory and visual information critical to pheromone-mediated optomotor anemotaxis. Furthermore, connections within SIP suggest another possible site for multimodal visual/olfactory convergence. SIP houses many arborizations from mushroom body output neurons (MBON) associated with the VL (described in *Drosophila*, Aso et al., 2014) (Ito et al., 2014). MBONs connect with the intrinsic MB neurons, Kenyon cells, that receive inputs from m-*mc*PNs. The projections of d-lALT *lc*PNs within SIP thus suggests indirect interactions between the mALT and lALT pathways. Additionally, loose fibers extend from the anterior optic tubercle, itself a large optic glomerulus, that receives input from optic lobe neuropils (medulla and lobula) into SIP although it is unclear whether these neurons actually terminate in SIP (Ito et al., 2014). Hence, similar to ΔLP, in SIP there is a potential confluence of olfactory information from the pathways segregated at the antennal lobe level in close proximity to potential visual inputs. The possible relationships between different pathways are represented schematically in Figure 13. Another factor in considering integration of signals in the protocerebrum be they olfactory-olfactory or olfactory-visual relates to the transmission speed along the various pathways to terminal synapses. The lALT pathway to the ΔLP is considerably shorter than that taken by PNs projecting through the mALT. It is conceivable that putative inhibitory inputs from l-*lc*PNs arrive at target neurons in advance of excitatory inputs from m-*mc*PNs and this may have a significant effect on the modulation of activity in these higher centers.

**Figure 13.**
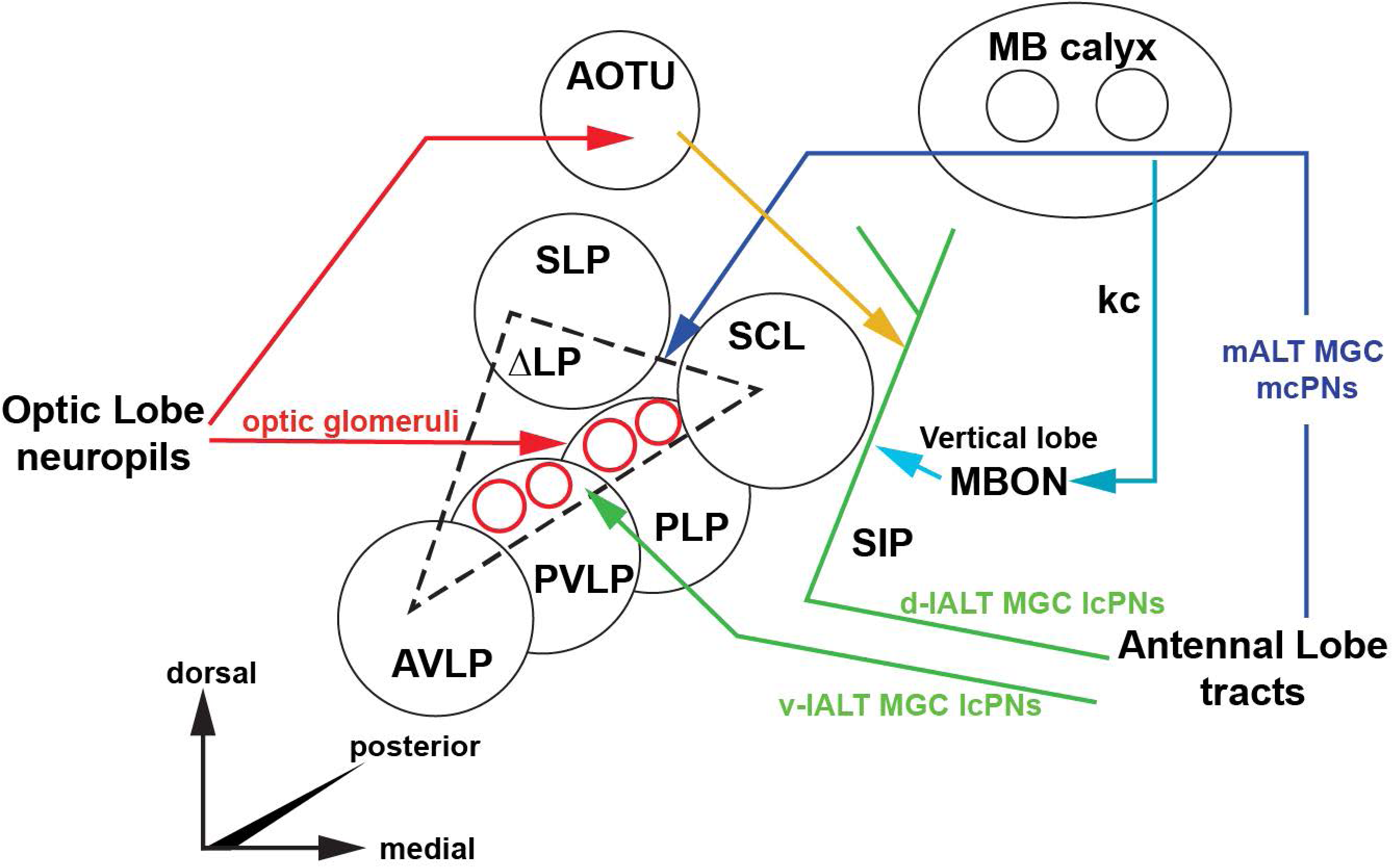
Schematic diagram showing a suggested relationship between olfactory (pheromone) and visual pathways converging in the ΔLP region of the moth protocerebrum. Based on current results and supported by other proposed models (Baker and Hansson, 2016). Projections from optic lobe neuropils target optic glomeruli in the dorsal regions of PVLP and PLP, overlapping with olfactory v-lALT MGC *lc*PNs. d-lALT MGC lcPNs might interact with mushroom body output neurons (MBON) that originate in the vertical lobe of the mushroom body (MB) where they receive inputs from the intrinsic MB kenyon cells (kc). There may also be interactions in this region with output neurons from the anterior optic tubercule (AOTU), itself an enlarged optical glomerulus. Thus, there appear to be multiple opportunities for convergence of olfactory input from different pathways onto target protocerebral neurons including between mALT PN and v-ALT PN (directly) and mALT PN and d-ALT PNs (indirectly through the MB). In addition, there is evidence for intermingling of visual and olfactory pathways through protocerebral optic glomeruli and v-ALT, d-ALT and mALT. Not shown: overlapping mlALT output in AVLP, PVLP areas that intersect with ΔLP. Connection between d-lALT through SIP and back into postero-dorsal ΔLP alongside the Posterior Lateral Fascicle. Multiple ipsilateral and contralateral connections of neuron shown in Fig. 9E.

The results reported here underscore the importance of including behaviorally relevant blends alongside their constitutive single odorants presented as short-duration, pulsatile stimuli in characterizing olfactory neurons since these parameters capture the odor stimulus conditions under which many insects respond behaviorally and reveal characteristics of neuronal response that may otherwise be concealed. Furthermore, because *mc*PNs and *lc*PNs are often excited by stimulation with single odorants their responses can appear similar making these two neuronal types difficult to distinguish solely on the basis of single odorant neurophysiological criteria. Since canonical uniglomerular m-*mc*PNs have been the predominant focus of many previous studies in moths, it seems likely that the contribution of *lc*PN output to higher centers has been underestimated.

## Acknowledgements

This project was supported by a grant from the National Science Foundation IOS-1147233 to NJV. Part of this study was conducted in partial fulfillment of the Ph.D dissertation by CFC in the Program in Neuroscience at the University of Utah.

## Author Contributions

Study conceptual framework (S-GL, CFC and NJV), experiments (S-GL, CFC, CK, JS and NJV), imaging (S-GL, CFC, CK, JS and NJV), data analysis (S-GL, CFC and NJV), manuscript preparation (CFC and NJV).

